# Bagging Improves Reproducibility of Functional Parcellation of the Human Brain

**DOI:** 10.1101/343392

**Authors:** Aki Nikolaidis, Anibal Solon Heinsfeld, Ting Xu, Pierre Bellec, Joshua Vogelstein, Michael Milham

## Abstract

Increasing the reproducibility of neuroimaging measurement addresses a central impediment to the advancement of human neuroscience and its clinical applications. Recent efforts demonstrating variance in functional brain organization within and between individuals shows a need for improving reproducibility of functional parcellations without long scan times. We apply bootstrap aggregation, or bagging, to the problem of improving reproducibility in functional parcellation. We use two large datasets to demonstrate that compared to a standard clustering framework, bagging improves the reproducibility and test-retest reliability of both cortical and subcortical functional parcellations across a range of sites, scanners, samples, scan lengths, clustering algorithms, and clustering parameters (e.g., number of clusters, spatial constraints). With as little as six minutes of scan time, bagging creates more reproducible group and individual level parcellations than standard approaches with twice as much data. This suggests that regardless of the specific parcellation strategy employed, bagging may be a key method for improving functional parcellation and bringing functional neuroimaging-based measurement closer to clinical impact.

## 1. Introduction

Understanding the brain’s organization is a fundamental goal of human neuroscience (Glasser et al., 2016) and important for clinical impact (Fox & Greicius, 2010). Brain parcellations are distinct partitions in the brain that can be individual contiguous areas, or networks made up of closely interacting regions (Eickhoff, Yeo, & Genon, 2018). Parcellation of the human brain into functionally homogeneous areas has seen impressive advances in recent years on resting state and multimodal neuroimaging (Eickhoff et al., 2018), often through the use of global similarity based clustering (Bellec, Rosa-neto, Lyttelton, Benali, & Evans, 2010; Craddock, James, Holtzheimer, Hu, & Mayberg, 2012; Thirion, Varoquaux, Dohmatob, & Poline, 2014), multivariate decomposition (Beckmann, Mackay, Filippini, & Smith, 2009; Varoquaux, Gramfort, Pedregosa, Michel, & Thirion, 2011), or local gradient-based methods (Cohen et al., 2008; Glasser et al., 2016; Gordon et al., 2014; Nelson et al., 2010; Wig et al., 2014; Xu et al., 2016). Such network- and region-level parcellations have enabled a greater understanding of fundamental attributes of the brain’s multi-level network structure (Bellec et al., 2015; Buckner, Krienen, Castellanos, Diaz, & Yeo, 2011; Margulies et al., 2016; Meunier, Lambiotte, Fornito, Ersche, & Bullmore, 2009; Yeo et al., 2011). Parcellations of the brain also commonly form the nodal backbone of network neuroscience studies (Hagmann et al., 2007; Zalesky et al., 2010), which have been successful in exploring the network structure of the brain and successfully predicting a wide variety of cognitive (IQ: (Finn et al., 2015; Shehzad et al., 2014); Complex Task performance: (Nikolaidis, Goatz, & Smaragdis, 2015)), developmental (Age: (Dosenbach et al., 2010; Liem et al., 2017)), physical (Aerobic fitness: (Talukdar et al., 2017)), and clinical phenotypes (Autism: (Abraham et al., 2017; Hong, Valk, Di Martino, Milham, & Bernhardt, 2018); Tourette’s Syndrome: (Greene et al., 2016)), and are an essential area of research for understanding human brain organization. Unfortunately, studies suggest that the amount of scan time required to achieve desirable levels of reproducibility and reliability appear to surpass the scan time in most clinical and cognitive neuroscience datasets already collected or currently underway (Laumann et al. 2015; Xu et al. 2016; Zuo et al. 2019; Greene et al. 2019). As such, there is also an increasing need for methods that can optimize the value of resting state acquisitions with insufficient data. The importance of functional parcellations in understanding brain organization and clinical, cognitive, and developmental phenotypes makes it critically important to assess new methods for improving their reproducibility and reliability.

Bootstrap aggregation, a popular technique from statistics and machine learning may assist in improving the reproducibility of functional parcellations. Bootstrapping and bootstrap aggregation are both general parametric or nonparametric procedures used to improve statistical inference along many dimensions, including improved estimates of certainty of an effect, estimating p-values, and accuracy. Bootstrapping refers to resampling a dataset with replacement, and has found a wide range of applications in science and statistics (Efron & Tibshirani, 1994). For example, bootstrapping is used for estimating the accuracy of a sample estimate and is based on the notion that a sample’s population can be modeled by resampling. Building on this concept, bootstrap aggregation, or bagging, was originally designed to resample data for prediction, and to aggregate the predictions across bootstrap samples to improve predictive performance (Breiman, 1996). Bagging is thought to reduce variability in the estimation through averaging (prediction labels or cluster membership) over multiple resampled datasets (Dudoit & Fridlyand, 2003). More recently, bagging has become an important technique for ensemble clustering, which is a way of creating a final clustering by aggregating a range of cluster solutions (Jia et al. 2011; Li and Ding 2008; Fischer and Buhmann 2003; Zhou 2012; Hong et al. 2009; Leisch 1999). In bagging for clustering, cluster partitions across all bootstrap samples form a cluster stability matrix by recording for each pair of observations the proportion of times they were clustered together in the bootstrap clusters (Dudoit and Fridlyand 2003; Fred and Lourenço 2008). This stability matrix is then used as input to a clustering procedure to create the final clustering. In other words, the aggregation of cluster solutions themselves become the features for ensemble clustering (Strehl & Ghosh, 2002).

Numerous studies outside of the imaging community (Dudoit & Fridlyand, 2003; Jia et al., 2011; Li & Ding, 2008; Strehl & Ghosh, 2002; Zhou, 2012), and one within (Hoyos-Idrobo, Schwartz, Varoquaux, & Thirion, 2015), have found that both: (1) within-sample instability, and (2) sensitivity to noise inherent to the cluster optimization procedure, are significantly attenuated by cluster ensembles generated through bagging. These and other efforts have demonstrated the impact of bagging cluster ensembles in improving the cluster homogeneity, stability, and accuracy of clusters (Boongoen and Iam-On 2018; Lange et al. 2004). We hypothesized that bagging cluster ensembles may specifically improve the between sample and between session reproducibility and reliability of individual and group parcellations as well. Given the importance of reproducibility of parcellations as outlined above, enhancing clustering procedures via bagging could dramatically improve reproducibility in a wide range of clinical and cognitive applications that rely on clustering.

In the present work, we leverage the Bootstrap Analysis of Stable Clusters (BASC) framework, and a recently developed Python extension, referred to as PyBASC. BASC is an implementation of bootstrap aggregation for functional neuroimaging analysis that was initially created to assess the stability of parcellations across a range of cluster resolutions and choose the ‘best’ resolutions that maximize within subject and within sample parcel quality and stability (Bellec et al., 2010). Such stability selection efforts have been demonstrated elsewhere to improve the estimation of structure in data (Meinshausen & Bühlmann, 2010) and are particularly well suited for choosing the number of clusters (von Luxburg, 2010). Initial efforts demonstrated that the circular block bootstrap (CBB) was advantageous for the estimation of robust associations between brain regions (Bellec, et al., 2008); later work focused on using CBB and bootstrapping at the group level to demonstrate the hierarchical nature of brain networks (Bellec et al., 2010, Bellec et al., 2015). PyBASC leverages the power of Nilearn (https://nilearn.github.io/), Scikit learn (Pedregosa et al., 2011), and Nipype (Gorgolewski et al., 2011) to build an open source Python package for BASC (GitHub: https://github.com/AkiNikolaidis/PyBASC; PyPI: https://pypi.org/project/PyBASC/). PyBASC allows both standard parcellation schemes (similar to Craddock et al. 2012), as well as bagging derived cluster ensembles as described above. Furthermore, we extend prior work in clustering and bagging ensembles by examining how bagged parcellations can be used to improve the reproducibility across sample, session, and study.

Keeping in line with extensive efforts at both cortical (Bellec et al., 2010; Craddock et al., 2012; Kelly et al., 2009; Kelly et al., 2012; Margulies et al., 2007) and subcortical parcellation (Barnes et al., 2010; Choi, Yeo, & Buckner, 2012; Garcia-Garcia et al., 2017; Janssen, Jylänki, Kessels, & van Gerven, 2015), we perform functional parcellation at multiple resolutions in both cortex and subcortex to assess the impact of bagging on reproducibility and reliability of these parcellations. While several cortical parcellations are well established (Glasser et al., 2016; Yeo et al., 2011), subcortex is an area where parcellations are less well-studied (Choi et al., 2012; Janssen et al., 2015). Subcortical regions are also in need of methods to improve reproducibility and reliability given that they show connectivity with notably lower reliability than the rest of the brain (Noble et al., 2017; O’Connor et al., 2017).

The goal of the present work is to improve the reproducibility of parcellations through the use of bagging enhanced ensemble clusters with PyBASC. We approach the issue of bagging for enhancing functional parcellation reproducibility across a range of parameters such as scan length, cluster number, sample size, cluster parameters, clustering method, and region of interest. We created parcellations of both cortex and subcortical structures using both bagging and a standard non-bagged approach leveraging two large datasets with multiple resting state acquisitions per participant (i.e., the Hangzhou Normal University (HNU1) from the Consortium for Reliability and Reproducibility (Zuo et al., 2014), and the Brain Genomics Superstruct Project (GSP) datasets (Holmes et al., 2015); see Table 1). We compare the similarity across scans, across individuals, and across groups to evaluate standard and bagging based parcellations in their reproducibility, reliability, and individual-to-group similarity. Importantly, reproducibility is quantified in different contexts: (1) of individual level parcellations within a sample, (2) of individual and group level parcellations between samples and sessions, and (3) of parcellations between group and individual level parcellations. To accomplish this, we use the definitional framework outlined in Table 2 and Section 2.5 & 2.6. We were motivated by the COBIDAS report (Nichols et al., 2017) and broader outlines for the establishment of reproducibility in metrology and biomedical science (JCGM, 2008).

**Table 1:** Overview of datasets used in the current study. HNU1-Hangzhou Normal University. GSP-Genomics Superstruct Project.

| Data Source | Sample Size | Data Length | Age | Sex | Voxel Size | TR |
| --- | --- | --- | --- | --- | --- | --- |
| HNU1 | 30 | 10x 10-Minute | 24.37, +/- 2.37 | 50/50 % F/M | 3.4, 3.4, 3.4 mm | 2000ms |
| GSP | 300 | 2x 6-Minute | 23.67, +/- 2.15 | 50/50 % F/M | 3.0, 3.0, 3.0 mm | 3000ms |

**Table 2.** Group-Level Analyses. Level of Generalization depicts which type of generalizability we tested with each analysis, see Methods sections 2.5-2.6. Columns show which kind of parameters would be kept constant (=) or changed (x) across analyses in the current study.

| Level of Generalization | Participants |  | MRI |  | Data Used | Figures |
| --- | --- | --- | --- | --- | --- | --- |
|  | Pop. | Sample | Scanner | Data |  |  |
| Between Scan | = | = | = | X | GSP 10 groups TRT | 7 |
| Between Session | = | = | = | X | HNU Reference & Replication | 8, 9, 10, 11, 12, 13 |
| Reliability | = | = | = | X | GSP 10 groups TRT<br>HNU Reference & Replication | 14, 15, 16 |
| Within Study<br>Between Sample | = | X | X | X | GSP 10 groups | 2 |
| Between Study<br>Between Sample | X | X | X | X | GSP 300<br>HNU Replication | 3, 4, 5, 6 |

We find that bagging has a significant impact on the reproducibility of parcellations at both the group and individual level, improving reproducibility even more than doubling scan time. Bagging enhanced parcellations are also less individually variable, exemplified by (1) large decreases in both within and between individual variance of individual level parcellations, and (2) increases in both univariate and multivariate measures of reliability. We also find that bagging increases individual-to-group similarity of the parcellations, meaning group level parcellations become more representative of the individuals in the group. <u>Bagging improves the generalizability, reproducibility, and reliability of both cortical and subcortical functional parcellations across a range of sites, scanners, samples, scan lengths, and clustering parameters.</u> Importantly, we find this to be true regardless of the specific cluster analysis algorithm being used. These results suggest that bagging may be an important component for achieving more robust parcellations in functional neuroimaging.

## 2. Materials and Methods

### 2.1 Overview

We aimed to approach a full-scale assessment of the impact of bagging on between sample reproducibility, between session reproducibility, and between session reliability. Towards this end, we apply PyBASC, a multi-level bagging approach for functional parcellation (See Methods Section 2.4 on PyBASC).

### 2.2 HNU1 and GSP Data

We assessed the impact of bagging on reproducibility of functional parcellation using the HNU1 (Zuo et al., 2014), and GSP datasets (Holmes et al., 2015). In the HNU1 dataset, 10-minute resting state scans were acquired in 30 young adults every three days for a month, for a total of 10 sessions per person. To test between session reproducibility and reliability we split the even and odd numbered sessions into 50-minute reference and replication datasets. From the 300 participants in the GSP, we created 10 groups of 30 age- and gender-matched participants to assess within study/between sample reproducibility. Using all 300 GSP participants combined, we also created a reference parcellation and compared it to the HNU1 replication dataset to assess the impact of bagging on between study/between sample reproducibility.

### 2.3 MRI Data

#### 2.3.1 MRI Acquisition

*HNU1. Anatomical*: 3D SPGR images were acquired on a GE 3T scanner (General Electric, Milwaukee, WI) with an 8 Channel head coil. Flip angle: 8 Degrees. TI: 450ms, TE: 60ms, TR: 8.06s, 180 slices, slice thickness: 1mm. Acquisition time: 5:01. *Rest*: T2* BOLD EPI sequence, GE 3T, 8 Channel head coil. Flip angle: 90 Degrees. TE: 30ms, TR: 2000ms, 43 slices, slice thickness: 3.4 mm. Acquisition time: 10:00.

*GSP Anatomical:* T1 MEMPRAGE images were acquired on Siemens 3T Magnetom Tim Trio scanner (Siemens Medical Solutions, Erlangen, Germany) with a 12 channel head coil. Flip angle: 7 degrees. TI: 1.1s, TE:1.5 / 3.4 / 5.2 / 7.0. TR 2.2s. 144 slices, slice thickness: 1.2mm. Acquisition time: 2:12. *Rest*: T2* BOLD epfid2d1_72 sequence, Siemens 3T Magnetom Tim Trio, 12 channel head coil. Flip angle: 85 degrees. TE: 30ms, TR: 3.0s, 47 slices, slice thickness: 3mm. Acquisition time: 6:12.

#### 2.3.2 Preprocessing

*Anatomical:* 1) AFNI skull stripping. 2) MNI 152 2mm template anatomical nonlinear registration with FNIRT. 3) Data transformed to 3mm. *Functional:* 1) Transformed to 3 mm. 2) Nuisance regression applied (white matter, CSF, global signal, motion, linear, and quadratic components of motion). 3) Friston 24-Parameter model used for volume realignment. 4) Nuisance band pass filtering between 0.01 and 0.1 Hz. 5) Spatial smoothing applied 6mm FWHM. Preprocessing in GSP and HNU1 structural and rest data were identical with the addition of de-spiking to the GSP resting state data. (HNU1-CPAC Version 1.0; GSP-CPAC Version 1.2).

### 2.4 PyBASC Parcellation

We first apply clustering in the same manner as is commonly applied in functional parcellation methods when not using bagging (See Figure 1; (Craddock et al., 2012; van den Heuvel, Mandl, & Pol, 2008). 1) We start with preprocessed functional MRI data; 2) this data is transformed from a voxel representation to a supervoxel representation through feature reduction such as Ward’s criterion based hierarchical agglomerative clustering; 3) time series for each supervoxel is extracted, and a correlation matrix is calculated, 4) clustering is applied to extract a specific number of clusters (K) for each individual’s correlation matrix and an adjacency matrix is created, and 5) individual level adjacency matrices are averaged together and 6) clustered again to reveal the group level clustering.

**Figure 1.**
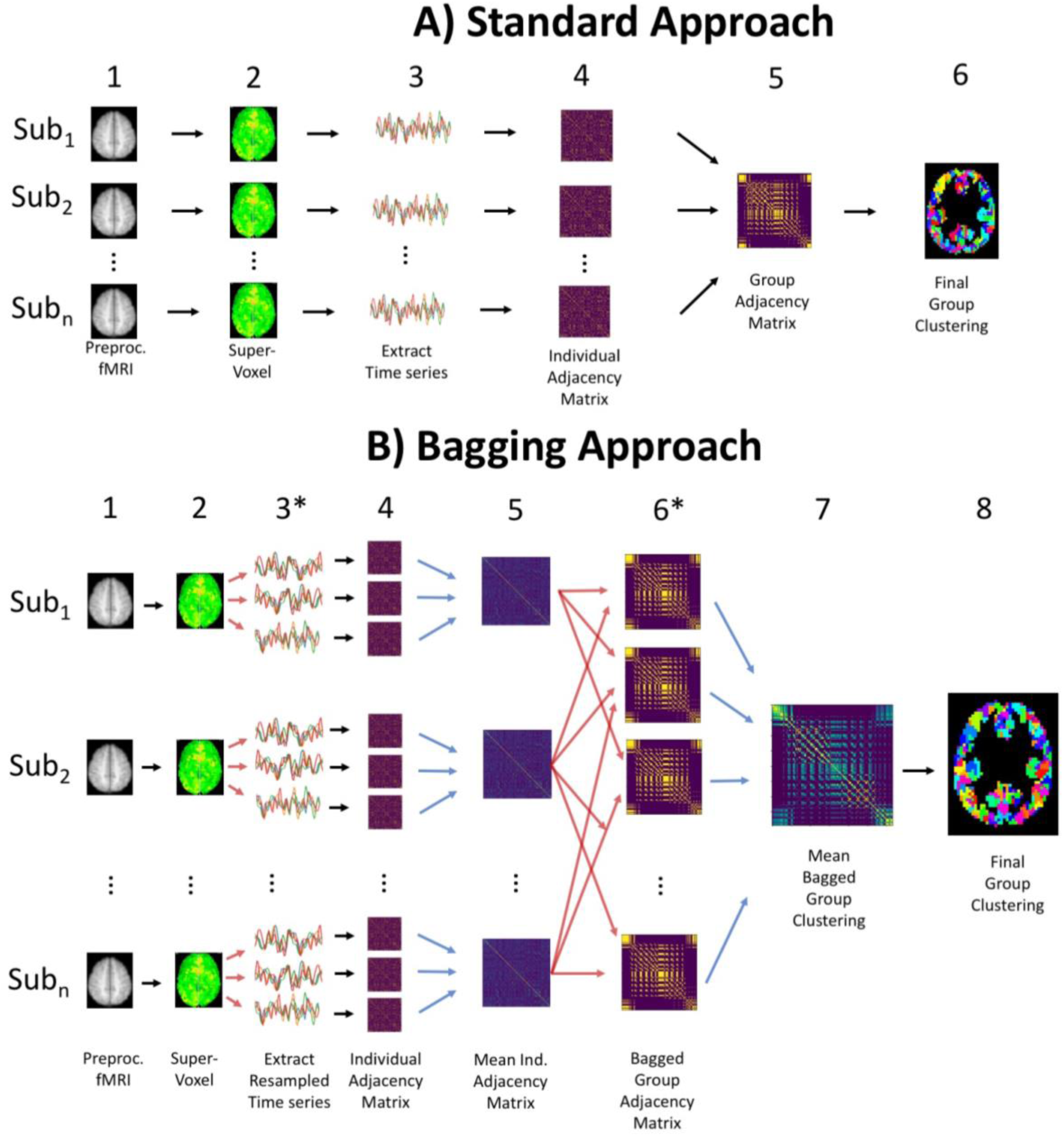
Parcellation Overview. A) Outline of different steps in standard clustering approach. B) Creating cluster ensembles through bagging follows the same overall structure as A, with the addition of individual level time-series resampling at step 3, and bagging across individuals in step 6. Red arrows show when bootstrap resampling occurs to create multiple resampled individual (3) and group level data (6). Blue arrows show when bootstrap aggregation happens to create averaged individual (5) and group level data (7).

This approach can be extended by adding bagging both within each individual’s time series and across individuals to create resampled groups, as first implemented by BASC (Bellec et al. 2010; Figure 1B). We explain this process briefly here (See Supplementary Materials 5.1). At the individual level, bagging is applied using a circular block bootstrap (CBB; window size = sqrt(number of time steps)). CBB has advantages of robustness for time series modeling and resampling compared to commonly used autoregressive models (Adolf, Baecke, Kahle, Bernarding, & Kropf, 2011; Adolf et al., 2014; Bellec, Marrelec, & Benali, 2008; Dupré la Tour et al., 2017; Liégeois, Laumann, Snyder, Zhou, & Yeo, 2017). We used this method on the individual time series to create many resampled 4D functional scans for each individual, and clustering was applied to each individual resampled 4D scan, resulting in cluster solutions for each individual level bootstrap. These cluster solutions are transformed to an adjacency matrix and are averaged together to create a mean adjacency matrix for each participant. We then bagged across individuals to create resampled groups (6), which are then averaged to create the group mean adjacency matrix (7) that is clustered to create the final parcellation (8). This multilevel bagging approach has been previously applied in the original BASC implementation in Octave and has been explained in greater detail elsewhere (Bellec et al., 2010; Garcia-Garcia et al., 2017); See Supplementary Materials 5.1 for more details about PyBASC). Here we have created a python-based implementation of this framework, PyBASC, to conduct the following reproducibility and reliability assessments. Overall, PyBASC and BASC follow the same algorithmic structure, however there are a few differences worth noting for the application here: 1) Dimension reduction is conducted through a region growing algorithm in BASC, whereas in PyBASC Ward’s criterion hierarchical clustering is applied. 2) Dimension reduction is applied at the group level in BASC, though can be at the individual or group level in PyBASC. Here we apply individual level dimension reduction for subcortical parcellation and group level dimension reduction for cortical parcellation to reduce computational load.

In the current work, we applied parcellation to both cortical and subcortical regions across a range of number of clusters (Subcortical K = 10, 18; Cortical K = 7, 20, 197, 444). We chose to apply bagging to the problem of both subcortical and cortical parcellation due to the computational advantage of the relatively few voxels present in our subcortical masks, which enables us to assess the impact of bagging across a wider range of parcellation parameters at a much lower computational cost than a full cortical parcellation. We chose K = 10, and 18 for the subcortical parcellation given our work and other literature has repeatedly shown these parcellations fit best for the striatum and thalamus (Behrens et al., 2003; Choi et al., 2012; Garcia-Garcia et al., 2017; Johansen-Berg et al., 2005). We chose K = 7, 20, 197, and 444 following previous analysis exploring multi-scale parcellation with BASC (Bellec, 2013; Bellec et al., 2010).

Functional parcellations often are created with strong spatial constraints to create more homogenous parcel shapes with smoother parcel boundaries (Craddock et al., 2012; Schaefer et al., 2018; Xu et al 2016). In this way such constraints can be considered a form of spatial regularization for the cluster optimization procedure, thus reducing variance by increasing bias and therefore increasing the reproducibility of the clustering outcomes. To provide the best test of the impact of bagging on parcellation reproducibility we use minimal spatial constraints in the current approach, enabling parcels that are discontinuous and even bilaterally homologous. While such an approach yields parcels with noisier boundaries than would be the case with stronger spatial constraints, such as Craddock et al. 2012, our approach allows the parcellation a better fit to a given sample and provides a stricter and more direct test of the impact of bagging. Notably, we also assessed the impact of bagging across spatial constraints (Supplementary Figure 5.2.4).

### 2.5 Reproducibility

#### 2.5.1 Concept

The definitions of concepts of reproducibility and replicability are numerous and often conflicting (Barba, 2018). We chose to follow the Joint Committee for Guides in Metrology (JCGM). This group is comprised of eight international organizations specializing in measurement, including the International Standards Organization (ISO), which the COBIDAS report follows closely (Nichols et al., 2017). According to the JCGM, Reproducibility is the similarity between the results of measurements of the same or similar measurands carried out under changed conditions of measurement (JCGM, 2008).

#### 2.5.2 Data Requirements

The changed conditions may include the method of measurement, observer, measuring instrument, reference standard, location, conditions of use, and time. Notably, statements of reproducibility must include a description of the specific conditions changed. In other words, when discussing reproducibility, it’s important to be specific about what is being changed and across which dimension(s) measurements are being reproduced. Reproducibility across sessions but not samples requires multiple assessments of the same sample and is known as between session reproducibility. Similarly, reproducibility across investigators requires different investigators but can use the same method and data, which is often referred to as computational reproducibility (Millman, Brett, Barnowski, & Poline, 2018). We extend this formal definition of reproducibility to include reproducibility across samples, which we call between sample reproducibility.

To support our tests of between sample, between session, and between scan reproducibility, we created several reference and replication datasets using the HNU1 and GSP datasets. In the HNU1 dataset, we created a test-retest dataset by combining the ten, 10-minute sessions into two five-session, 50-minute scans. Our reference and replication datasets were created through a concatenation of each participant’s even and odd numbered session’s respectively. To assess the interaction of bagging and length of acquired data on reproducibility, we performed parcellation across the first 3, 5, 10, 15, 20, 25, and 50 minutes of the replication data. We split the 300 participants of the GSP data into 10 groups of 30 participants that were age and sex matched to the HNU1 dataset, and we used these 10 groups to assess both between sample and between scan reproducibility, given these two 6-minute scans were acquired in the same session. We also concatenated both 6-minute sessions and used all 300 participants of the GSP dataset to create a large reference parcellation, and we compared this reference to the replication sets of the HNU1 data to test between sample reproducibility.

#### 2.5.3 Testing Methods

To measure between session reproducibility, we assessed the similarity between the reference and each of the replication datasets using spatial correlation of these group level mean adjacency matrices, and the adjusted Rand index (ARI) of the group level cluster labels. Given that bagged clustering is non-deterministic, we assessed the variance in our parcellations by repeating our parcellations 20 times for each replication dataset across each amount of scan time and bootstrap aggregation for the subcortical parcellation. For the cortical parcellation we repeated the parcellations 10 times, with the HNU1 10-minute replication data. Since the standard, non-bagged, clustering approach is deterministic, we needed a method to assess the variance in between session reproducibility of the parcellations. To overcome this issue, we created 20 new datasets for each amount of the scan lengths by resampling the 50-minute replication dataset (See Supplementary Materials 5.1 for details on this resampling method). We applied PyBASC without bagging across each of these 20 datasets to estimate the variance in between session reproducibility for the standard parcellation condition.

We assessed the effect of bagging on within study/between sample reproducibility of functional parcellation by comparing the parcellations between 10 age and sex-matched groups of 30 participants from the GSP dataset. Since we had 10 groups, we did not need to resample and repeat our bagged or standard parcellations as above. Each of the two 6-minute scans were used for each group, as well as a concatenated 12-minute scan. We compared the similarity of group level parcellations between the 10 groups for each scan length, and we compared the parcellations of the ten groups for both 6-minute scans. Second, we assessed the effect of bagging on between study/between sample reproducibility. We used the same 12-minute scans from all 300 GSP participants together to create a reference parcellation. For assessing cortical parcellation reproducibility, we compared the GSP reference to the 10-minute HNU1 data. For assessing subcortical parcellation reproducibility we assessed the GSP parcellation against a range of HNU1 scan lengths.

### 2.6 Reliability & Discriminability

#### 2.6.1 Concept

Reliability is a metric of the inter- and intra-individual stability of a measurement across multiple occasions (Zuo & Xing, 2014), and can be measured with different indices. Intraclass correlation (ICC) is a descriptive statistic that relates the within individual to between individual variance to get an indication of univariate reliability (Shrout & Fleiss, 1979). Discriminability is a multivariate metric of reliability that takes the full set of features across all observations into account to compute a multivariate index of reliability (Bridgeford et al. 2020; https://github.com/neurodata/r-mgc).

#### 2.6.2 Data Requirements

Calculating test retest reliability requires at least two observations of the same individuals in the same group. In the current study, we compute ICC using two measurements from each individual in both the HNU1 and GSP samples.

#### 2.6.3 Testing Methods

We assessed whether bagging would improve both the univariate and multivariate reliability of our functional parcellations, and how scan length would have an impact on these results. Using the same set of functional parcellations across multiple datasets calculated in Section 2.5 Reproducibility, namely the reference and multi-length replication datasets, we calculated the ICC for each cell in the individual level mean adjacency matrix data, and discriminability for the individual mean adjacency matrix as a whole.

## 3. Results

### 3.1 Bagging Improves Between Sample Reproducibility

Improving between sample reproducibility is key for creating generalizable scientific discovery. We assess two different levels of between sample reproducibility here: within study and between study, and we hypothesized that bagging should have a beneficial impact on both levels. As a first step, we investigated the effect of bagging on the within study/between sample reproducibility of our subcortical parcellations (Figure 2). We split the 300 GSP participants into 10 groups of 30 and computed the correlation of the subcortical parcellations group mean adjacency matrices (Figure 2 A, B), and the ARI of their cluster labels between groups (Figure 2 C, D). We found that while increasing the time of the scan from 6 to 12 minutes significantly improved reproducibility when measured by both correlation and ARI (K = 10; Correlation: Kruskal-Wallis = 58.52, df = 1, p-value < 0.0001; ARI: Kruskal-Wallis = 10.79, df = 1, p-value < 0.005), we found that the impact of bagging improved reproducibility more than increasing the scan length (K = 10; Correlation: Kruskal-Wallis = 401.26, df = 1, p-value < 0.0001; ARI: Kruskal-Wallis = 278.9, df = 1, p-value < 0.0001), indicating that the 6 minute scans could become more reproducible than scans twice as long with the use of bagging. We also found that bagging significantly decreased the variance in the reproducibility estimates for both 6 minute and 12 minute scans, but had a greater impact on the 12 minute scans (6 min ARI variance 0 vs 400 bootstraps: F(267,267) = 0.716, p-value < 0.01; 12 min ARI variance 0 vs 400 bootstraps F(89,89) = 1.6712, p-value < 0.05).

**Figure 2.**
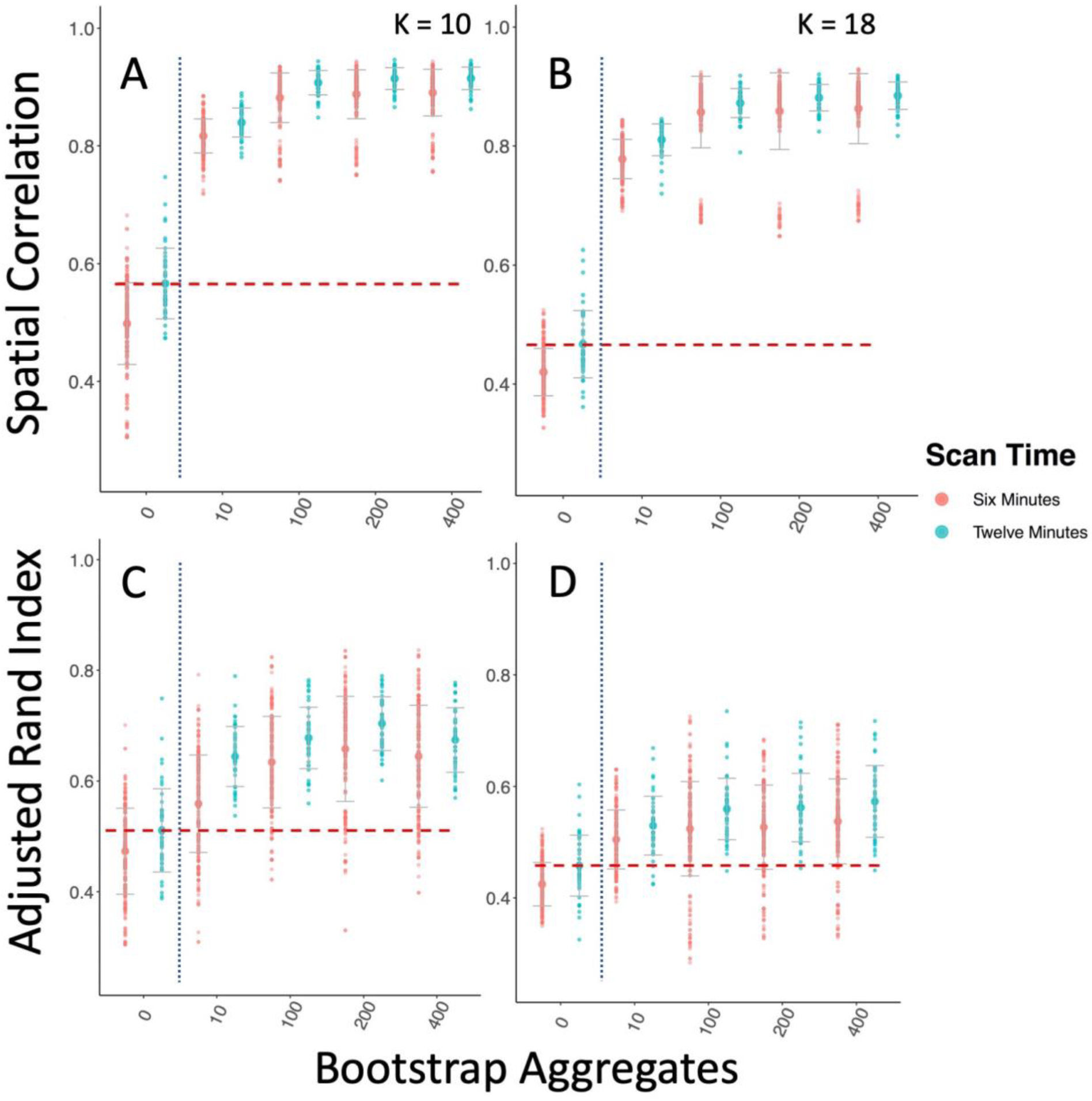
Bagging Improves Reproducibility of Group Level Within Study/Between Sample Subcortical Parcellation. We compare the parcellations from 10 groups of 30 participants from the GSP sample to one another. The vertical dotted blue line marks the standard clustering approach on the left, and the bagging approach on the right. The horizontal dashed red line shows the highest mean reproducibility of the standard parcellation approach. A & B Spatial correlation between mean group adjacency matrix of 10 GSP groups with 10 and 18 clusters. C & D ARI between cluster labels of 10 GSP groups with 10 and 18 clusters.

As a further test of generalizability, we tested between study/between sample reproducibility of both cortical parcellation between the GSP and HNU1 studies (Figure 3; Figure 5). These studies differ on many criteria including scanner type and population. While the GSP data was collected in the USA with Siemens 3T Magnetom Tim Trio scanners, the HNU1 data was collected in China with a GE 3T scanner. We hypothesized that bagging should not only improve the reproducibility of the parcellations, but also that it may have an impact on the consistency of the replication parcellations as well. We conducted a range of tests to assess these concepts. First, using the 300 individuals from the GSP study to create a reference cortical parcellation, we assessed the correlation of the group mean adjacency matrices (Figure 3 A), and the ARI of the group cluster labels (Figure 3 B). We demonstrated that cortical parcellations were more reproducible between studies using bagging; for example, for K = 444, bagging improves ARI (Kruskal-Wallis test comparing 400 vs 0 bootstraps = 12.79; p < 0.0005). We also found that bagging lowered the variance of ARI across repeated parcellations (Test of unequal variance of ARI 0 vs 400 bootstraps: F = 106.67, p < 0.0001). This shows that the parcellations of the replication dataset were not only more consistent with the reference dataset, but also that repeated parcellations of the replication dataset were more consistent with each other when bagging was used than when it was not. One of the challenges in parcellation is that as we increase spatial scale, we tend to lose reproducibility (Arslan et al., 2018), making obtaining highly detailed maps of the brain problematic. We find that bagged parcellations at higher resolution (K = 197) are as reproducible as network level parcellations without bagging (K = 7; Figure 3). This indicates that bagging also assists in achieving reproducible functional parcellations at higher resolutions.

**Figure 3.**
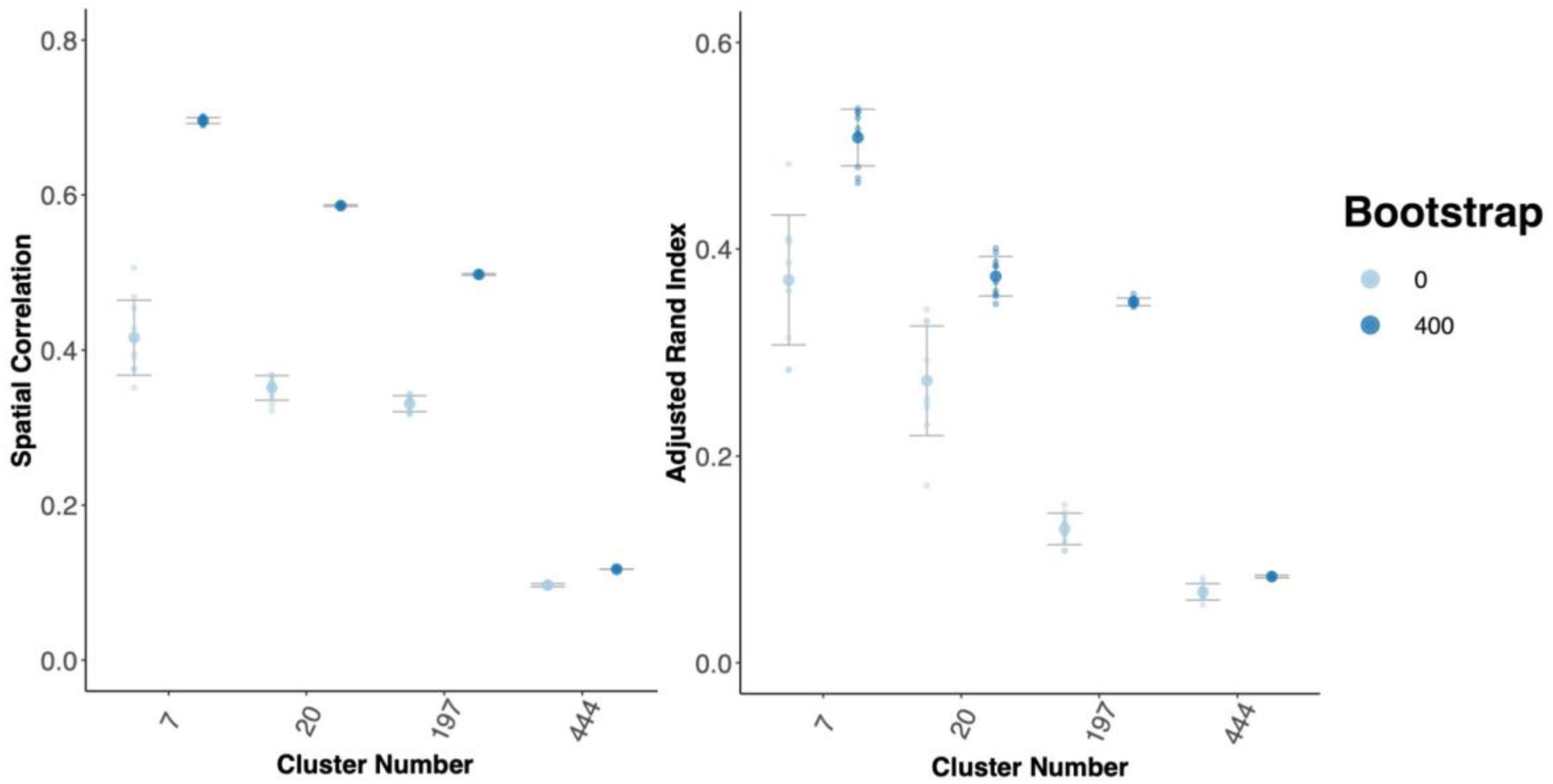
Bagging Improves Reproducibility of Group Level Between Study/Between Sample Cortical Parcellations. We compared a reference parcellation created from the 12-minute combined scan from 300 individuals from the GSP to 10 repeated parcellations of 10 minutes of HNU1 data. A) Spatial correlation between the group mean adjacency matrix for the GSP and HNU1 parcellations. B) ARI between the cluster labels for the GSP and HNU1 parcellations.

Third, we found that bagging had a significant impact on the between study/between sample reproducibility of subcortical parcellation, comparing the GSP reference to the HNU1 dataset (Figure 4; Figure 6). Both correlation and ARI of the parcellations improves with scan time from 10 to 50 minutes as expected. However, we expected that bagging would improve the parcellation labels equally for both the 10- and 50-minute replication samples. What we found instead, is that while both improved significantly from 0 to 400 bootstraps, the 10-minute HNU1 dataset improved much more than the 50-minute (K = 10, 10 minute: t = −13.087, df = 21.389, p-value < 0.0001; 50 minute: t = −3.2407, df = 21.75, p-value < 0.005). When visualized (Figure 6; Supplementary Material 5.2.1), we see this difference is largely due to the fact that both the GSP reference and the 10-minute HNU1 scans split the bilateral caudate into two clusters, whereas the 50-minute HNU1 parcellation did not. In fact, the 50-minute HNU1 replication and reference data yielded a seemingly better quality parcellation overall, with less variance on the edges between parcels, and more homotopic similarity, which can be expected given homotopic connectivity is known to be especially high (Zuo et al., 2010). We also visualized the voxelwise differences in parcellation labels on average over all HNU1 bootstrapped conditions and found most differences could be found in the boundary regions between parcels (See Supplementary Figure 5.2.1). Of note here, is that the clusters created have minimal spatial constraints applied to them, implying that neither the cluster solutions themselves, nor the improvement in reproducibility come from the clusters being forced into a similar spatially or anatomically constrained configuration (Shen, Tokoglu, Papademetris, & Constable, 2013). In fact, we see many clusters in the HNU1 50-minute Reference and Replication solution that are anatomically distinct but functionally united bilateral homologues (i.e. the left and right putamen cluster together, the left and right caudate cluster together, etc.).

**Figure 4.**
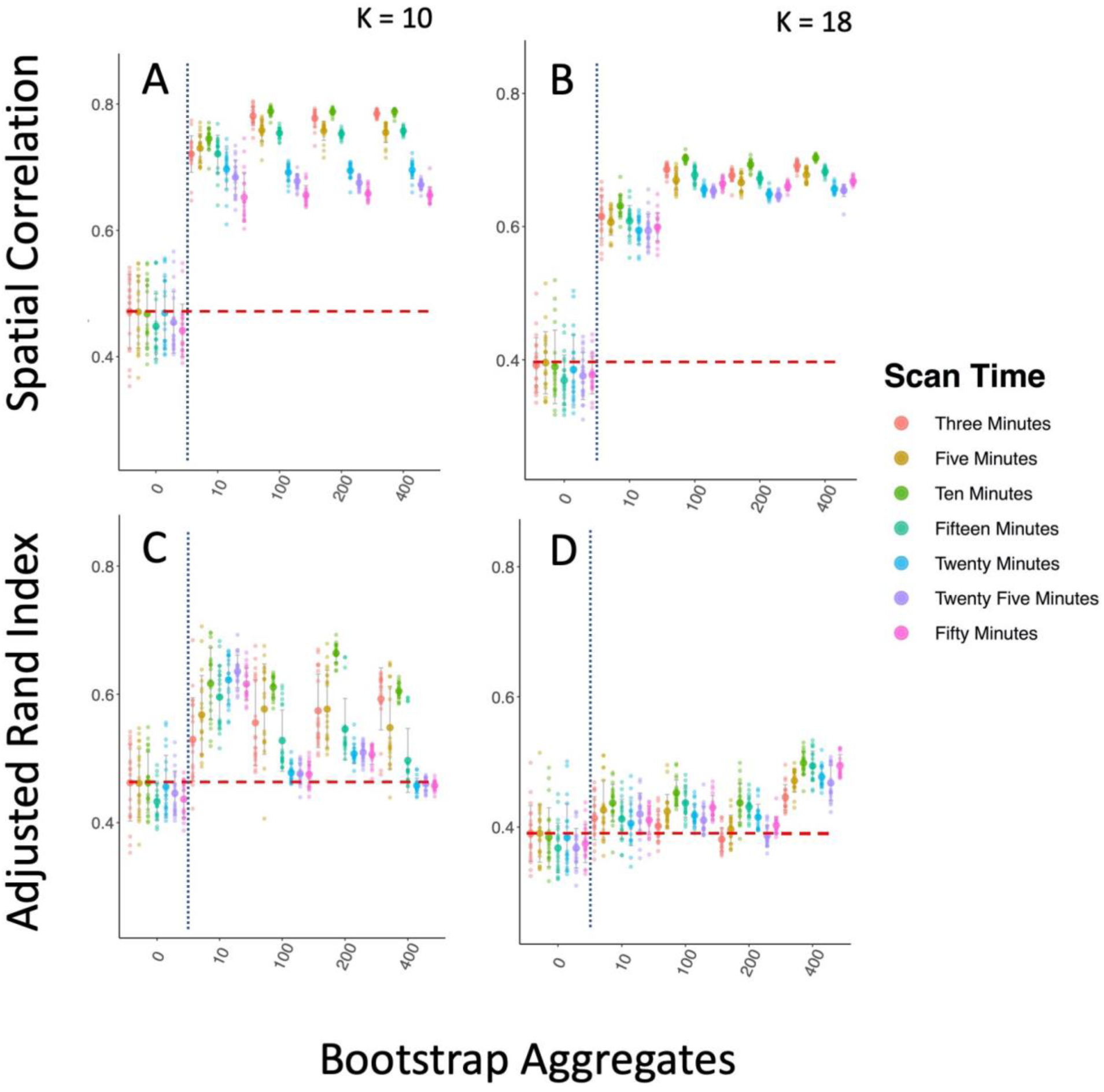
Bagging Improves Reproducibility of Group Level Between Sample Subcortical Parcellations. Bagging improves the reproducibility of subcortical brain parcellation between the GSP and HNU1 datasets, though results more mixed than the cortical results. We compared a reference parcellation created from the 12-minute combined scan from 300 individuals from the GSP to 10 repeated parcellations of 10 minutes of HNU1 data. The vertical dotted blue line marks the standard clustering approach on the left, and the bagging approach on the right. The horizontal dashed red line shows the highest mean reproducibility of the standard parcellation approach. A & B Show spatial correlation between the group mean adjacency matrix for the GSP and HNU1 parcellations for K = 10 and K = 18. C & D Show ARI between the cluster labels for the GSP and HNU1 parcellations for K = 10 and K = 18.

**Figure 5.**
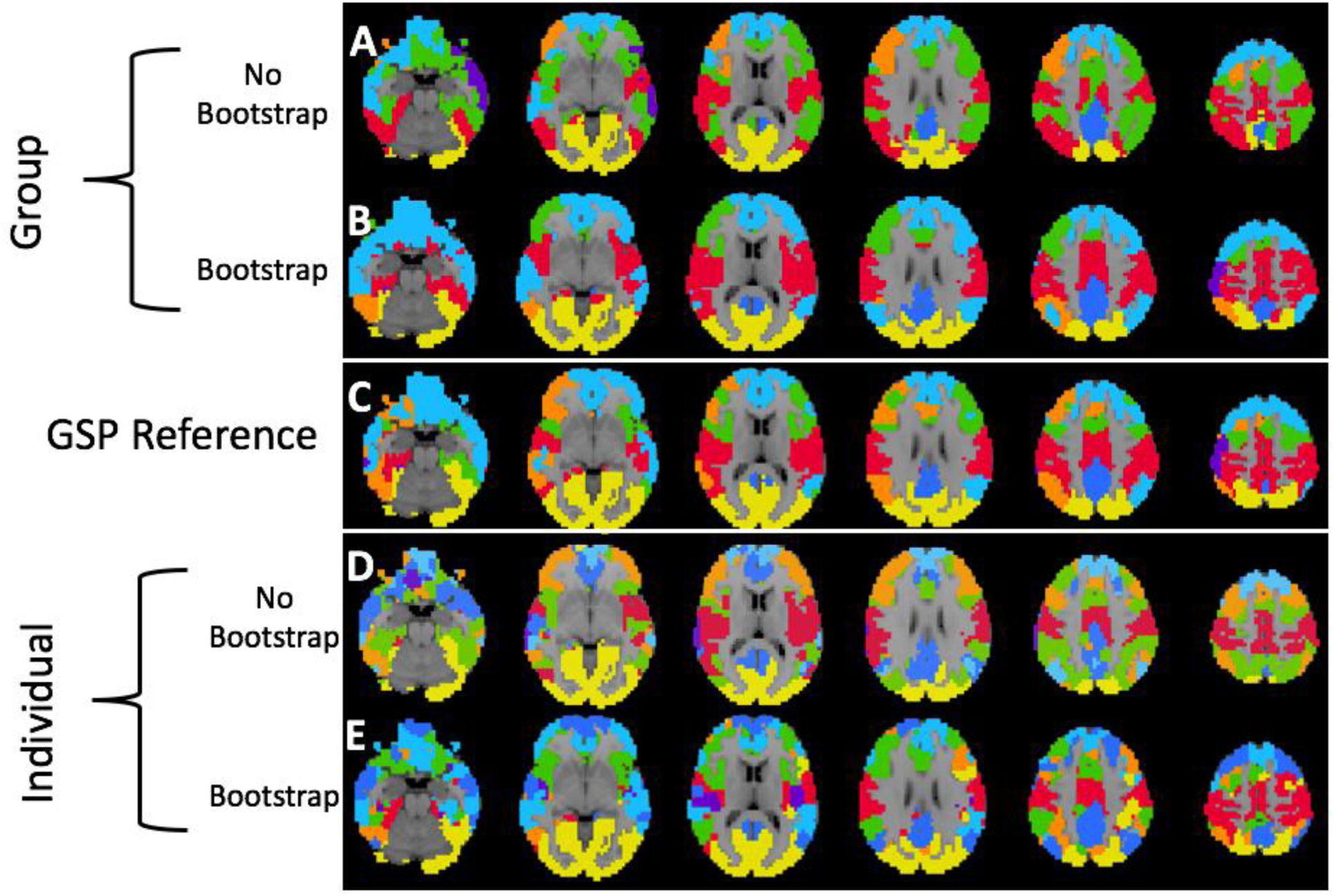
Visualization of Group and Individual Level Between Study Cortical Parcellations. We show the similarity of the HNU1 parcellation to the GSP reference parcellation with K = 7 clusters as shown in Figure 3. We use a K-nearest neighbor spatial filter on these parcellations to improve ease of visual comparison. Notably, both the group and individual level parcellations with 400 bootstraps are closer to the GSP reference parcellation than either of the standard parcellations. A) Group-level parcellation with standard approach, and B) bootstrapping. C) GSP Reference parcellation. D) Individual-level parcellation with standard approach, and E) bootstrapping.

**Figure 6.**
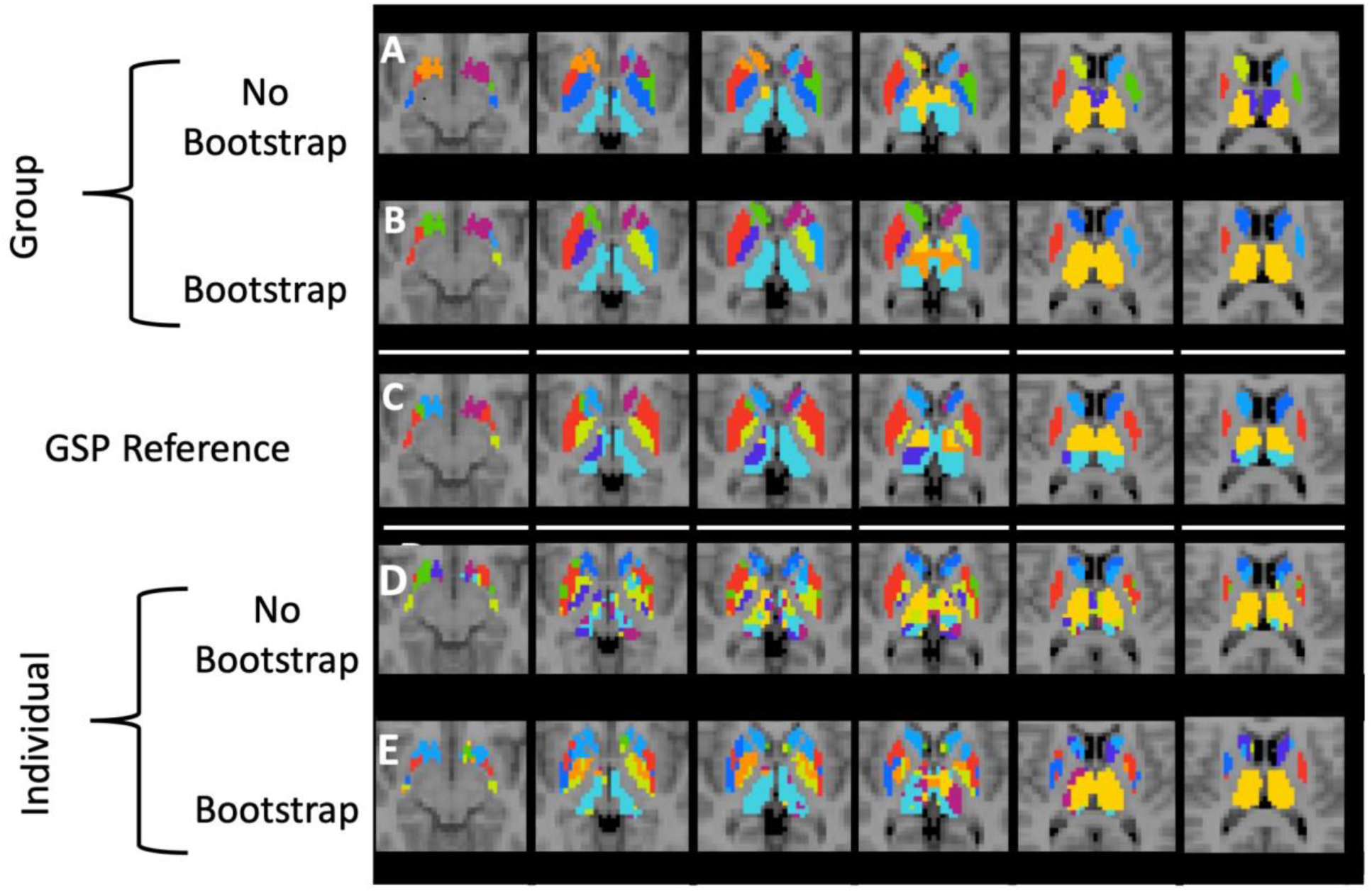
Visualization of Group and Individual Level Between Study Subcortical Parcellations. We show the similarity of the HNU1 parcellation to the subcortical GSP reference parcellation with 10 clusters as shown in Figure 4. Both the group and individual level bootstrapped parcellations are notably more similar to the GSP reference than either of the standard parcellation counterparts. A) Group-level parcellation with standard approach, and B) bootstrapping. C) GSP Reference parcellation. D) Individual-level parcellation with standard approach, and E) bootstrapping.

### 3.2 Bagging Improves Between Session and Between Scan Reproducibility

Whereas creating parcellations with high between sample reproducibility is key for generalizing scientific discovery, creating parcellations with higher between session and between scan reproducibility is key for decreasing uncertainty in measuring change over time, which is key for clinical and developmental neuroscience. We tested between scan reproducibility using the subcortical parcellations of the GSP dataset. We tested between session reproducibility using both subcortical and cortical parcellations of the HNU1 dataset.

First, we found that our subcortical parcellations of the GSP dataset, where we compared the effect of bagging on between scan reproducibility of 10 separate groups of 30 participants, had higher between scan reproducibility with bagging with as little as 6 minutes of data (Figure 7). This suggests that long scan times are not required for bagging to have a significantly beneficial impact on a parcellation’s between scan reproducibility of the group mean adjacency matrix and cluster labels (K = 10; Correlation: Kruskal-Wallis = 14.29, df = 1, p-value < 0.0005; ARI: Kruskal-Wallis = 10.57, df = 1, p-value < 0.005).

**Figure 7.**
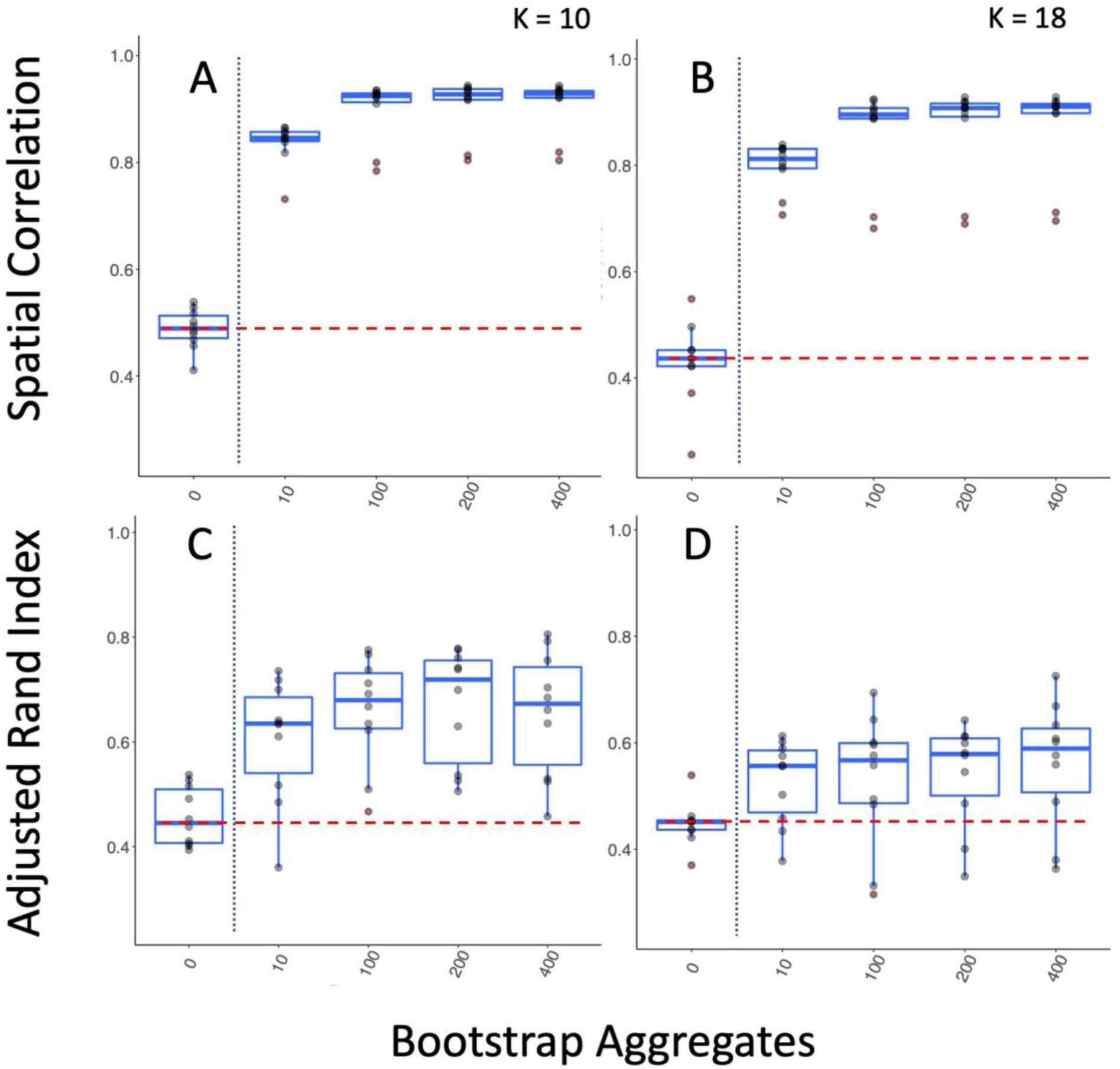
Bagging Improves Reproducibility of Group Level Within Sample/Between Scan Subcortical Parcellations. We split the 300 participants of the GSP dataset into 10 groups of 30 and compare the parcellations of their first and second functional scan to one another. The vertical dotted blue line marks the standard clustering approach on the left, and the bagging approach on the right. The horizontal dashed red line shows the highest mean reproducibility of the standard parcellation approach. A & B show the improvement in bagged parcellations measured by spatial correlation of the group mean adjacency matrix for 10 and 18 clusters respectively. C & D show the improvement in bagged parcellations measured by the ARI of the cluster labels for 10 and 18 clusters respectively.

We also found that our cortical parcellations had higher between session reproducibility with bagging in the HNU1 dataset (Figure 8; Figure 12). Comparing the K = 197 parcellation, we see significant improvements in both between session reproducibility of group mean adjacency matrices (Kruskal-Wallis = −78.86, df = 9.22, p-value = < 0.0001), and group cluster labels (Kruskal-Wallis = −33.30, df = 9.82, p-value < 0.0001). If parcellations have higher between session reproducibility on average but have high variance across repeated parcellations we might not have a good grasp on the extent to which a particular parcellation will match across time. Therefore, it’s also important to minimize the variance in the between session parcellation reproducibility. We found that bagging also significantly decreases the variance in between session reproducibility of the group mean adjacency matrix (F(9,9) = 83.46, p-value < 0.0001), suggesting that the bagged parcellations are not only more reproducible (higher mean ARI), but that this reproducibility is more stable as well (lower ARI variance). We also find that bagged parcellations at higher resolutions are more reproducible than network-level parcellations without bagging (K = 7; Figure 8). This replicates our earlier findings (Figure 3), that bagged parcellations enable robust investigations of brain organization at higher spatial scales.

**Figure 8.**
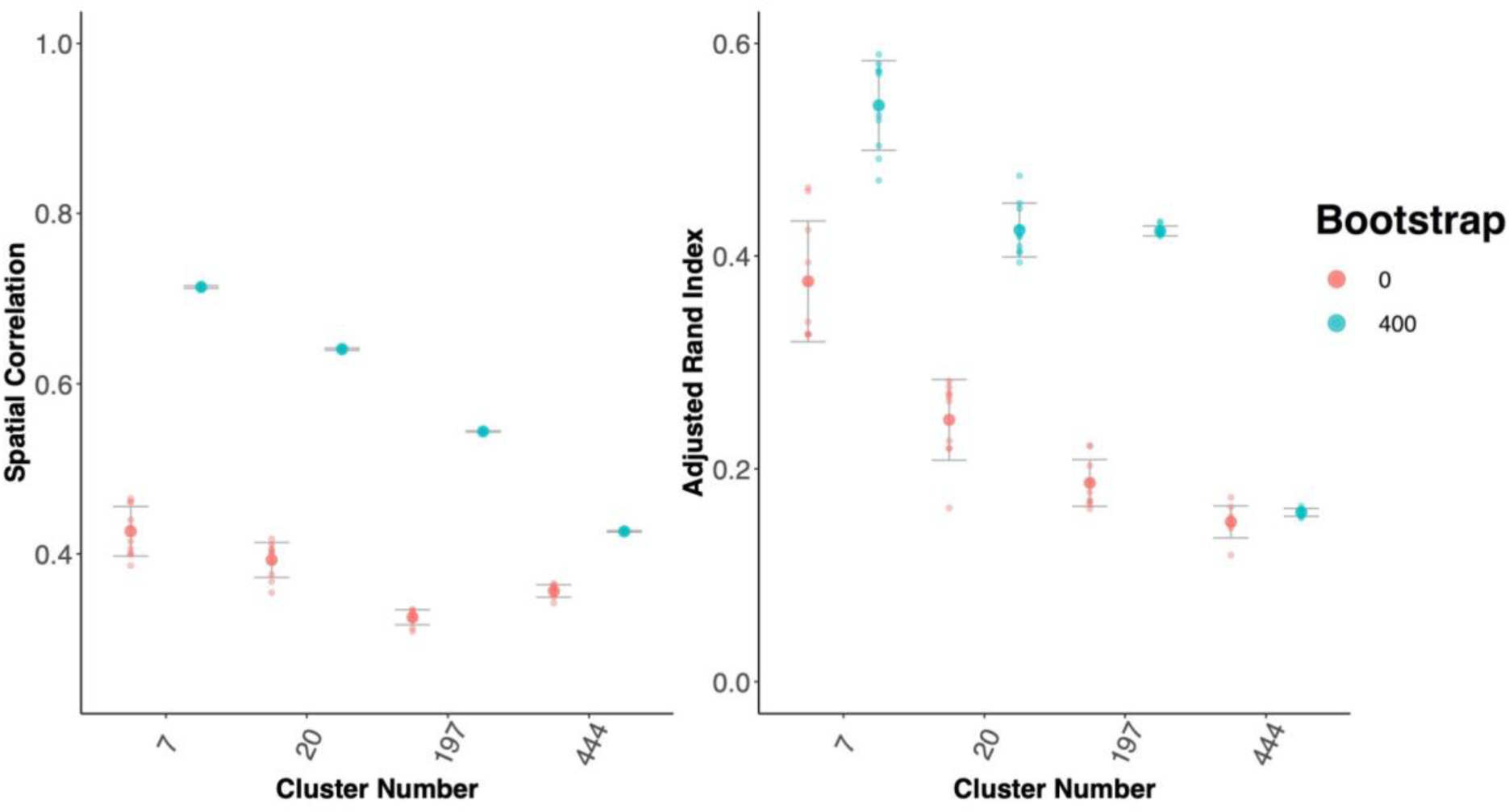
Bagging Improves Reproducibility of Group Level Between Session Cortical Parcellations. We compared the HNU1 reference to the HNU1 replication cortical parcellation. A) Spatial correlation between the group mean adjacency matrix for the reference and replication parcellations. B) ARI between the cluster labels for the reference and replication parcellations.

Finally, we tested the effect of bagging on the between session reproducibility of subcortical parcellations in the HNU1 dataset (Figure 9; Figure 13). We found that scan length had a highly significant impact on the between session reproducibility of parcellations. For instance, comparing 5- and 50-minute group mean adjacency matrix correlation and cluster label ARI with 0 bootstraps (K = 10; Correlation: Kruskal-Wallis = 14.14, df = 1, p-value < 0.0005; ARI: Kruskal-Wallis = 12.18, df = 1, p-value < 0.0005), and with 400 bootstraps (K = 10; Correlation: Kruskal-Wallis = 29.27, df = 1, p-value < 0.0001; ARI: Kruskal-Wallis = 29.27, df = 1, p-value < 0.0001). Notably, the impact of time was enhanced in the bagging condition, suggesting bagging and scan time can work synergistically to provide parcellations with high between session reproducibility. We also found that while between session reproducibility was improved with bagging, these reproducibility estimates for shorter scan times were not improved as was the case with longer datasets (10 minutes; K = 10; Correlation: Kruskal-Wallis = 29.27, df = 1, p-value <0.00001; ARI: Kruskal-Wallis = 1.00, df = 1, p-value > 0.05). We also visualized the voxelwise differences in parcellation labels on average over all HNU1 bootstrapped conditions and found most differences could be found in the boundary regions between parcels, along a dorsal-anterior ventral-posterior gradient (See Supplementary Materials 5.2.2), supporting the existence of small scale gradients in thalamic functional organization (Lambert, Simon, Colman, & Barrick, 2017).

**Figure 9.**
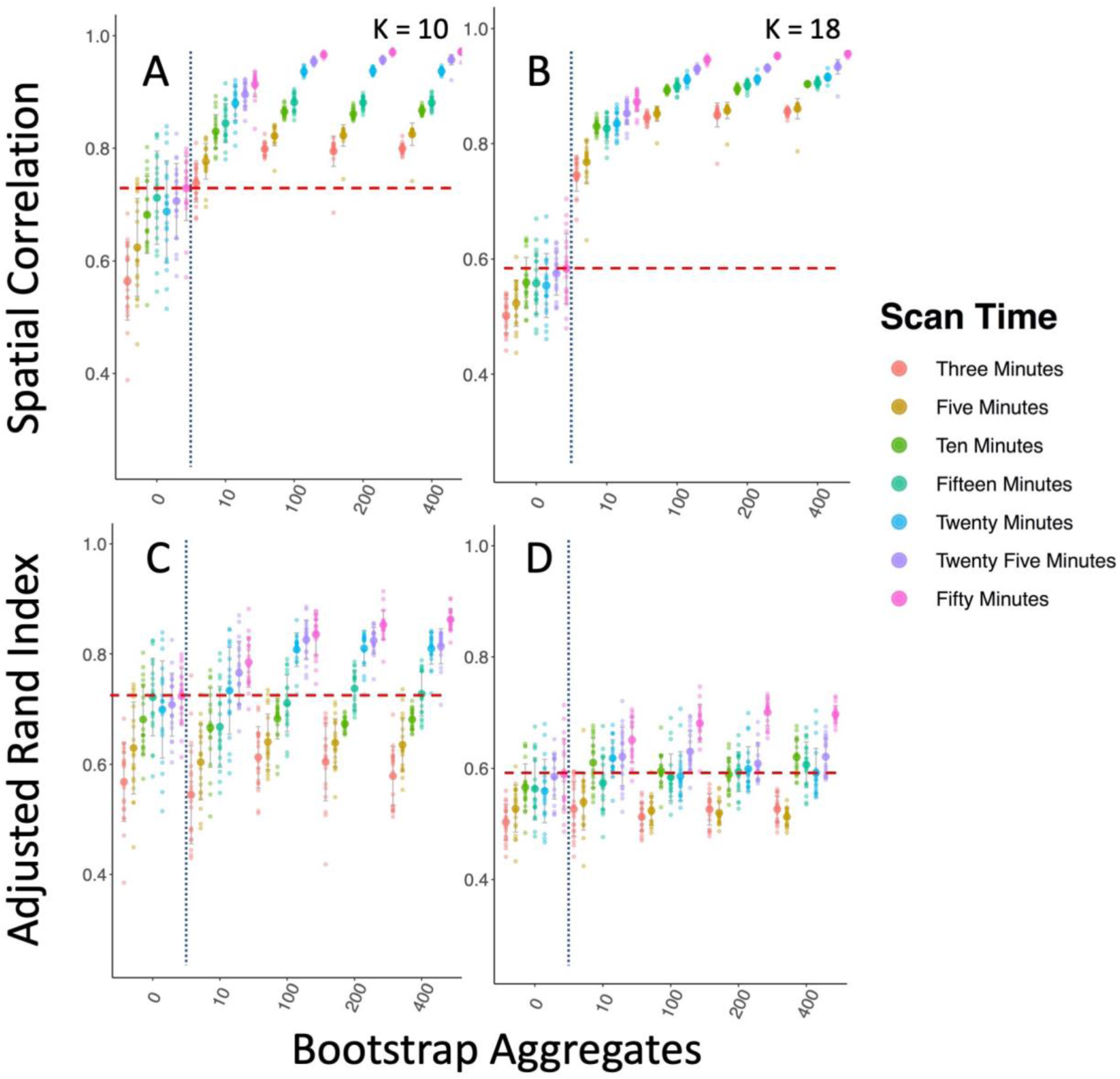
Bagging Improves Reproducibility of Group Level Between Session Subcortical Parcellations. We compared the HNU1 reference to the HNU1 replication subcortical parcellations across different lengths of time. The vertical dotted blue line marks the standard clustering approach on the left, and the bagging approach on the right. The horizontal dashed red line shows the highest mean reproducibility of the standard parcellation approach. A) Spatial correlation between the group mean adjacency matrix for the reference and replication parcellations. B) ARI between the cluster labels for the reference and replication parcellations.

We also see that the improvement in between session reproducibility from bagging is significantly greater than increasing scan length. For example, for the standard clustering condition, increasing the scan time from 20 minutes to 50 minutes does not yield a significant improvement in between session reproducibility of group mean adjacency matrix correlation or ARI (K = 10; Correlation: Kruskal-Wallis = 2.90, df = 1, p-value > 0.05; ARI: Kruskal-Wallis = 0.797, df = 1, p-value > 0.05), but increasing the bagging from 0 to 400 has a significant effect on the between session reproducibility of the 20-minute data (K = 10; Correlation: Kruskal-Wallis = 29.27, df = 1, p-value < 0.0001; ARI: Kruskal-Wallis = 17.58, df = 1, p-value < 0.0001). This demonstrates that bagging can improve reproducibility even more than doubling scan time, replicating our previous findings with the between-sample reproducibility in the GSP dataset (Figure 2). We also found that bagging had a significant impact on reducing the variance in between session reproducibility across multiple parcellations, replicating our results with the cortical parcellation (20 Minutes; K = 10; Correlation: F(19,19) = 311.14, p-value < 0.0001; ARI: F(19,19) = 9.7003, p-value < 0.0001). While the cortical parcellation demonstrated that only 10 minutes of data was needed for bagging to improve between session reproducibility in the HNU1 data (Figure 8), with the subcortical parcellation we saw that bagging did not improve the reproducibility for less than 20 minutes of data (Figure 9). This may be due to random elements of sampling those first 3, 5, 10, or 15-minute portions of the HNU1 dataset, or differences in the parcellation favored by lower amounts of data compared to higher amounts, as we saw in comparing the HNU1 50-minute parcellation to the GSP 12-minute reference parcellation in section 3.1.

We found that bootstrapping led to significant improvements in the reproducibility of individual level subcortical parcellations (Figure 11). Overall we saw that doubling scan time and using bootstraps led to significant improvements in reproducibility of individual level parcellations as measured by both spatial correlation (K = 10; Doubling scan time Kruskal-Wallis test effects between > 39 for all scan lengths; Using bootstraps Kruskal-Wallis test effects > 720 for all scan lengths) and ARI (Doubling scan time Kruskal-Wallis test effects > 26 for all scan lengths; Using bootstraps Kruskal-Wallis test > 99 for all scan lengths), and these improvements were consistent across all scan lengths (all p< 0.0005). Notably, we saw that bagging improved individual-level reproducibility more than doubling scan time as measured by spatial correlation for all time lengths, and as measured by ARI for scan lengths over 10 minutes. Furthermore, we replicated this effect in our cortical parcellation as well. We find that in our cortical parcellation, bagging significantly increases individual-level reproducibility as measured by spatial correlation of adjacency matrices and ARI of cluster labels across all cluster levels (all p < 0.00005; Kruskal-Wallis test: K = 7; Corr: 389.19, ARI: 200.55; K = 20; Corr: 433.73, ARI: 415.41, K = 197; Corr: 158.35, ARI: 449.25; K = 444; Corr: 86.48; ARI: 449.25). This is key for many approaches aiming to improve individual level parcellation (Gordon et al., 2017; Kong et al., 2019; Laumann et al., 2015; Xu et al., 2016).

Importantly, we also find that even bagging at the group level alone is enough to improve the between session reproducibility of the functional parcellations (Figure 10). This shows that individual level parcellations that are not bagging-enhanced can be bagged to improve the group level representation. This finding is important for other multi-level cluster estimation problems, such as diffusion weighted parcellation, which may benefit from the addition of group level bagging (Glasser et al., 2016; Klein et al., 2007; Mars et al., 2011).

**Figure 10.**
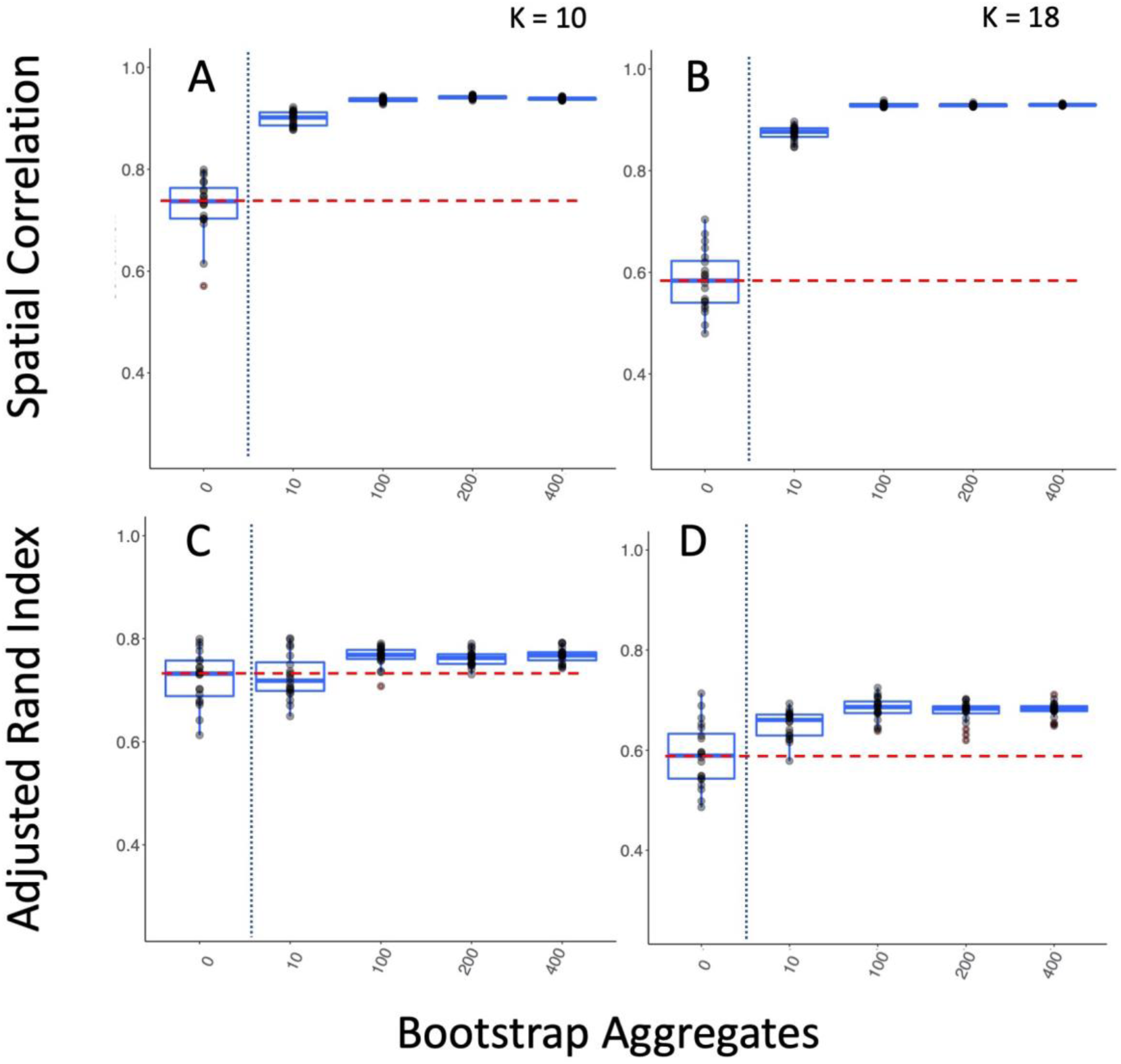
Group Bagging Alone Improves Reproducibility of Within Study/Between Session Subcortical Parcellations. We compare the subcortical parcellations of the 50-minute replication and reference datasets from HNU1 when only group level bagging is performed. The vertical dotted blue line marks the standard clustering approach on the left, and the bagging approach on the right. The horizontal dashed red line shows the highest mean reproducibility of the standard parcellation approach. A & B show the improvement in the spatial correlation of the group level mean adjacency matrix for K = 10 and K = 18, while C & D show the improvement in the ARI of the group cluster labels for K = 10 and K = 18.

**Figure 11.**
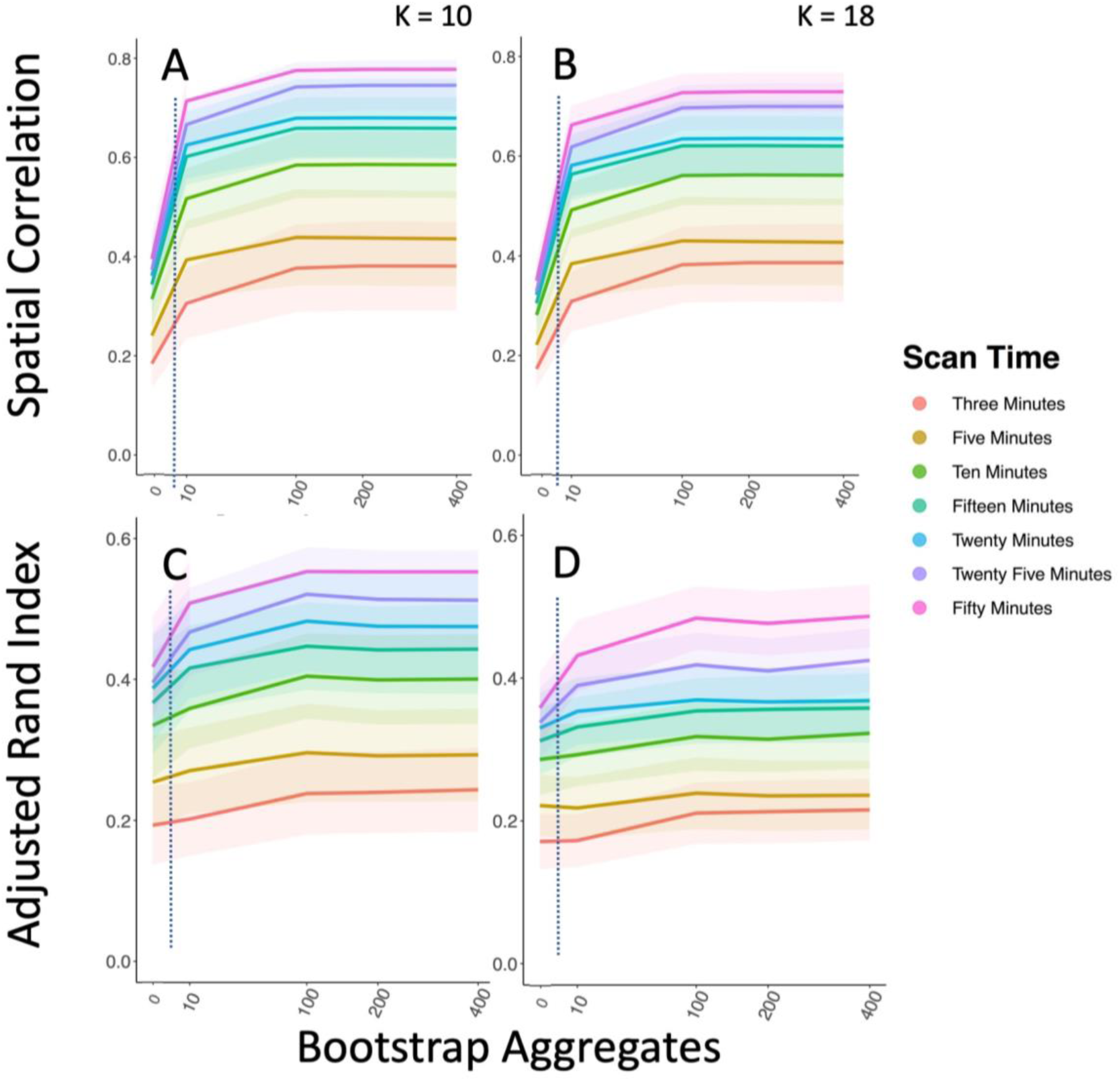
Bagging Increases reproducibility of Individual Level Parcellations. Individual-level bagging improves between session reproducibility of individual level parcellations across a range of bagging levels and range of scan times for the HNU1 data. We compare the individual mean stability matrix for each participant (A, B) and the individualized cluster labels (C, D).

**Figure 12.**
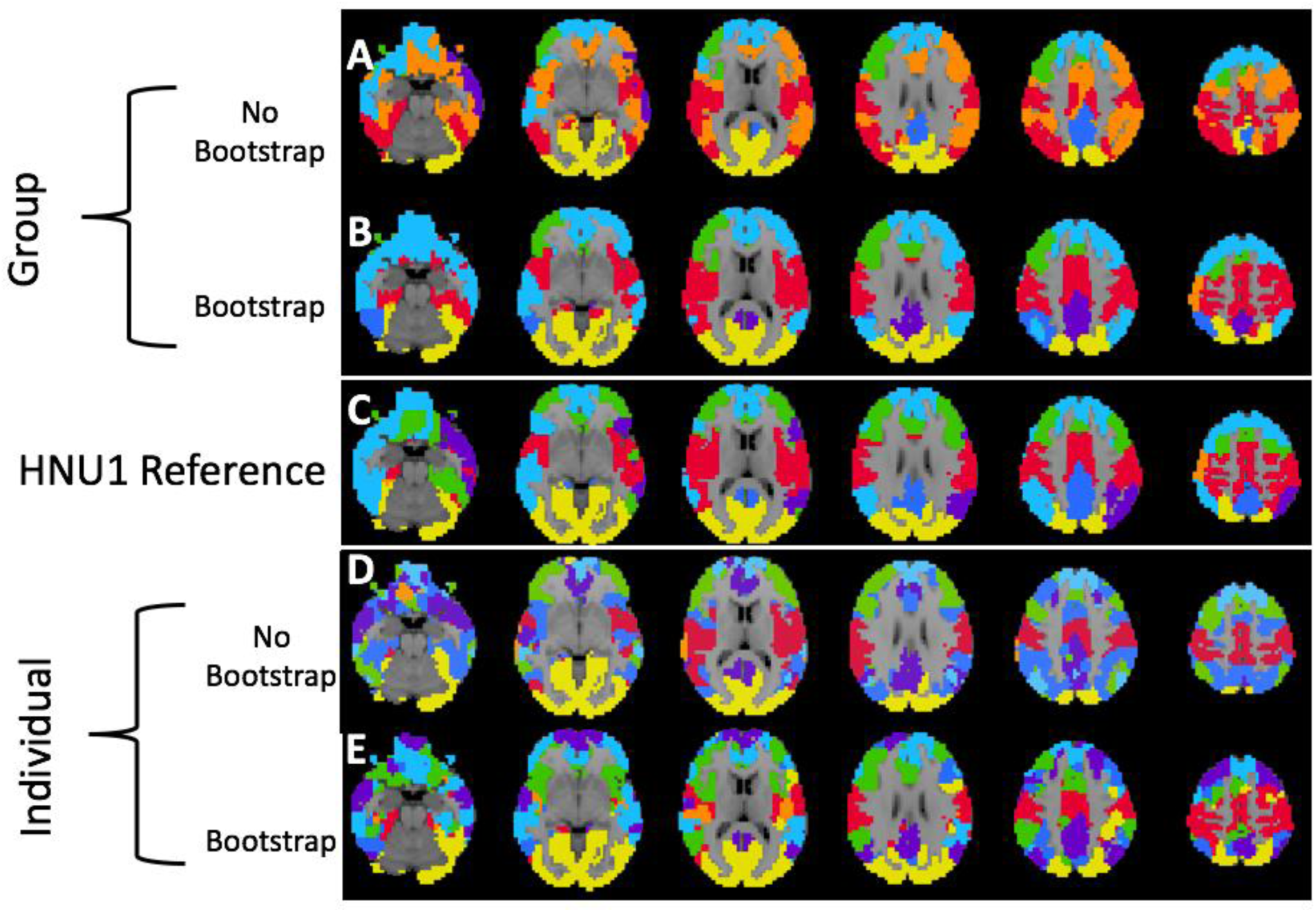
Visualization of Group and Individual Level Between Session Cortical Parcellations. We show the similarity of the HNU1 reference parcellation to the HNU1 replication parcellation with 7 clusters as shown in Figure 8. We use a K-nearest neighbor spatial filter on these parcellations to improve ease of visual comparison. Both the group and individual level bootstrapped parcellations are notably more similar to the HNU1 reference than either of the standard parcellation counterparts. We see notably more evidence for large-scale network similarities between the HNU1 reference and the bootstrapped parcellations at both the individual and group level, though the individual level is less obvious. A) Group-level parcellation with standard approach, and B) bootstrapping. C) HNU1 Reference parcellation. D) Individual-level parcellation with standard approach, and E) bootstrapping.

**Figure 13.**
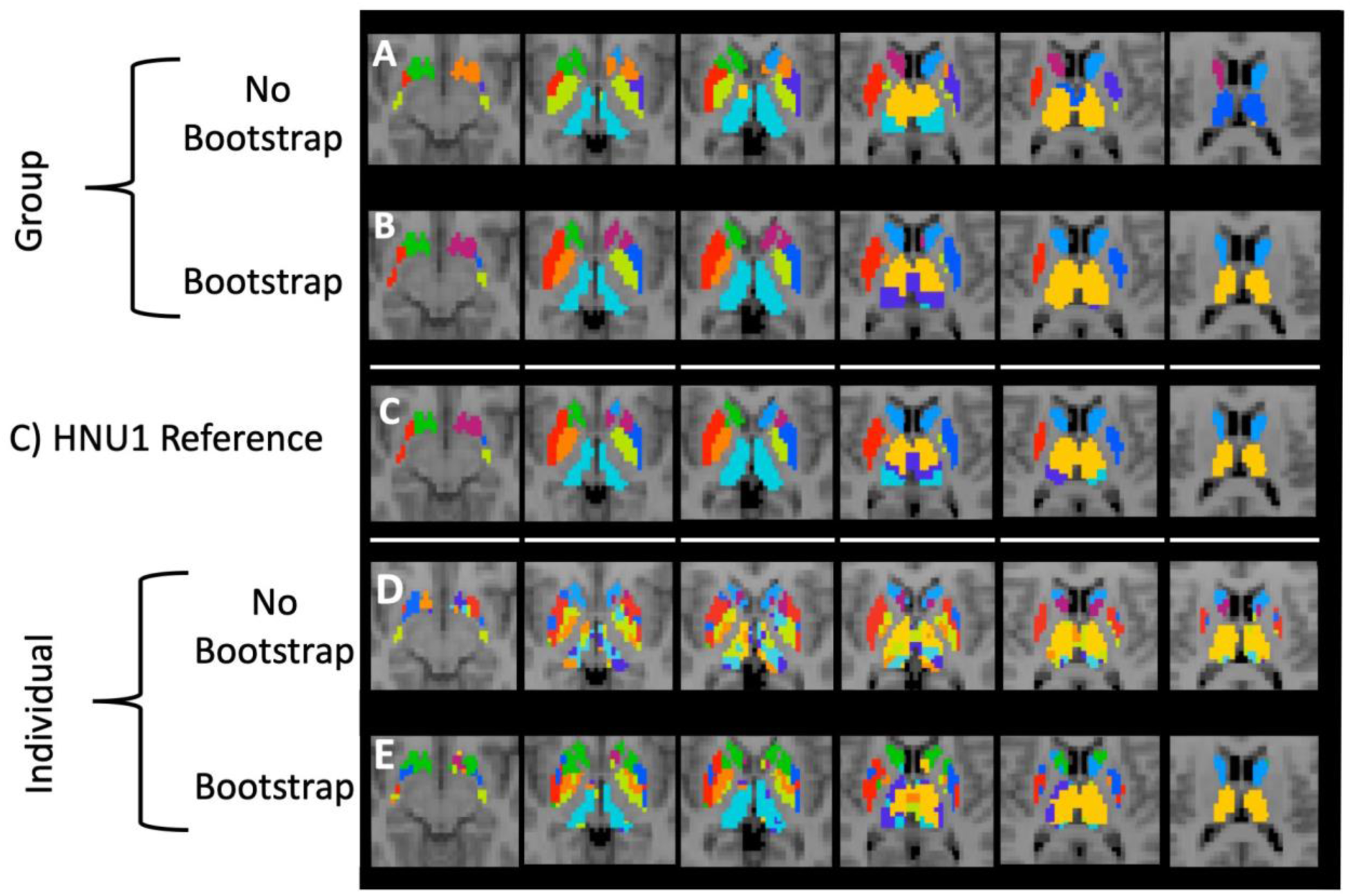
Visualization of Group and Individual Level Between Session Subcortical Parcellations. We show the similarity of the HNU1 reference parcellation to the subcortical HNU1 replication parcellation with 10 clusters, as shown in Figure 9. Both the group and individual level bootstrapped parcellations are notably more similar to the HNU1 reference than either of the standard parcellation counterparts. A) Group-level parcellation with standard approach, and B) bootstrapping. C) HNU1 Reference parcellation. D) Individual-level parcellation with standard approach, and E) bootstrapping.

Notably, we also tested the impact of spatial constraints on the effect of bagging on parcellations (Supplementary Results 5.2.3). Spatial constraints are an important consideration when creating any functional parcellation, without spatial constraints the clustering algorithms find groups of voxels with the most similar activation patterns regardless of their spatial distribution. The addition of spatial constraints moves the clustering algorithm to favor voxels that neighbor the cluster over others. If constraints are strong enough, anatomically disconnected parcels are not possible and the shape of parcels will be driven more by the spatial constraint than the actual underlying functional connectivity. Given that existing functional parcellation techniques use a wide range of spatial constraint approaches, we also varied the spatial constraints employed here to test if bagging would improve the reproducibility of parcellations across different spatial constraint criteria. We found that bagging improves subcortical parcellation between session reproducibility when considering fully spatially constrained clustering approaches as well (Subcortical; K = 10; Correlation: Kruskal-Wallis = 29.27, df = 1, p-value < 0.0001; ARI Kruskal-Wallis = 29.27, df = 1, p-value < 0.0001).

Finally, we also assessed the generalizability of the impact of bagging across functional parcellation techniques. We included an assessment of the impact of bagging on the between session reproducibility of subcortical parcels (K = 10) while using spectral clustering rather than Ward’s criterion based hierarchical agglomerative clustering (Supplementary Figure 5.2.5). Replicating our other results with Ward’s criterion based hierarchical agglomerative clustering (Figure 9), we found that bagging also improved the between session reproducibility using spectral clustering (Subcortical; 20 minutes; K = 10; Correlation: Kruskal-Wallis = 68.4, df = 4, p-value < 0.0005; ARI: Kruskal-Wallis = 10.48, df = 4, p-value < 0.05; 50 minutes; Correlation: Kruskal-Wallis = 85.72, df = 4, p-value < 0.0005; ARI: Kruskal-Wallis = 59.52, df = 4, p-value < 0.0005). This demonstrates that the bagging enhanced improvements in reproducibility generalize across clustering methods.

### 3.3 Bagging Improves Univariate and Multivariate Reliability

We found that bagging led to improvements in univariate estimates of reliability calculated with ICC of the voxel-wise mean adjacency matrices (Figure 14 A,B,C). Each of the entries in these matrices is the number of times across individual level bootstraps that two voxels are put into the same cluster. As such, the individual mean adjacency matrix is a representation of the average clustering solution across bootstrap aggregations for a given individual. ICC is calculated based on the relative size of variance between individuals compared to variance within individuals. We see that bagging led to large decreases in both between individual variance (MSR), and within individual variance (MSE; Figure 14). Notably, the decrease in MSE is slightly higher than the decrease in MSR, leading to a rise in ICC (Figure 14 D,E). In this case, ICC does not capture that bagging leads to a ten-fold decrease in variance, both within and between individuals’ individual level parcellation estimates.

**Figure 14.**
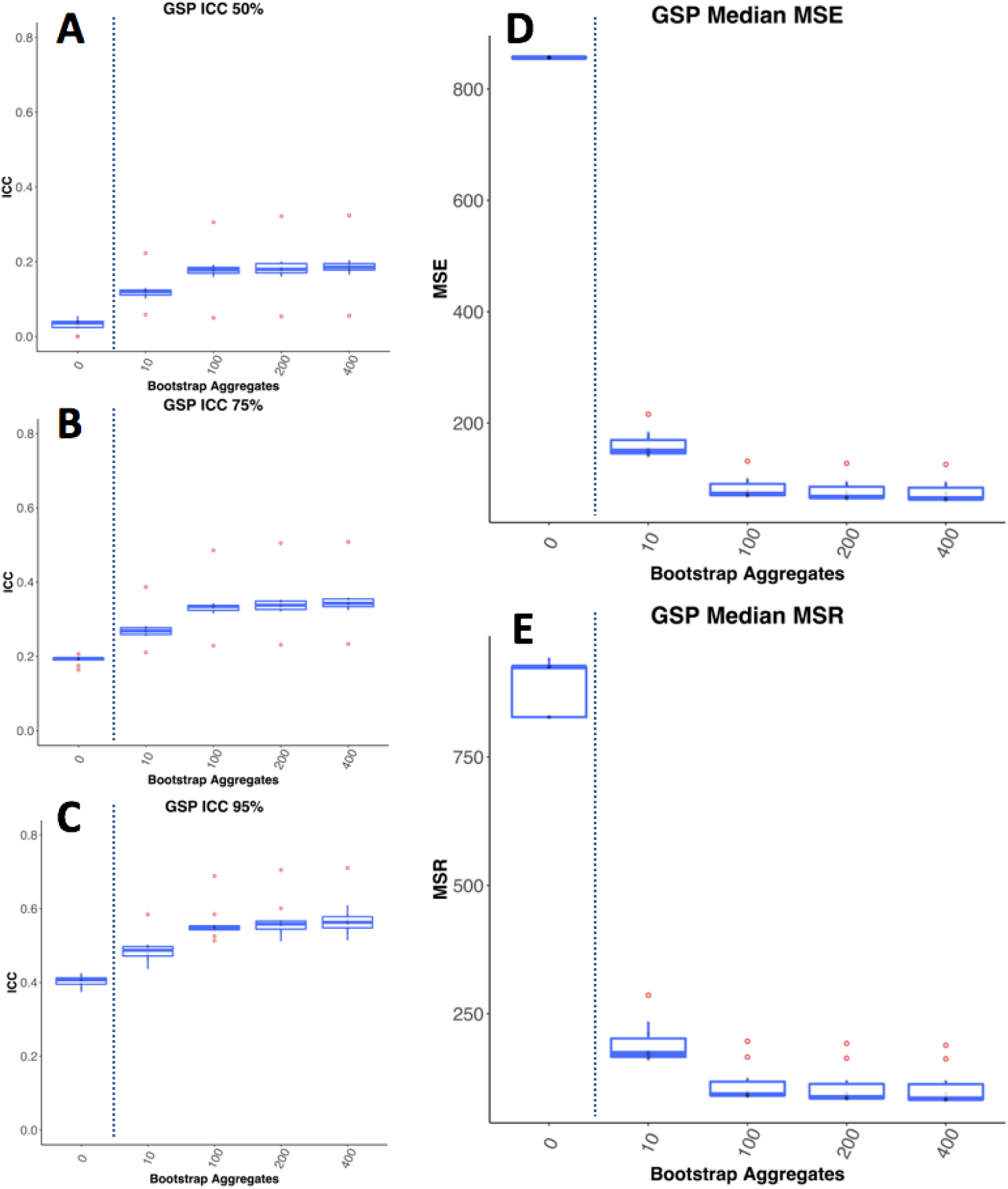
Bagging Improves ICC of Subcortical Parcellations. A, B, C show the top 50%, 75%, and 95% ICC values for the individual level mean adjacency matrices in the 10 GSP groups. D and E show the mean squared error within individuals and between individuals respectively. We see that bagging improves ICC. It also has a dramatic impact on both within individual variance (MSE) and between individual variance (MSR), with samples with the smallest drop in MSR showing the greatest pickup in ICC.

Motion can be a source of highly reproducible signal in MRI data and can highly influence estimates of reliability (Power, Barnes, Snyder, Schlaggar, & Petersen, 2012; Yan et al., 2013). We wanted to make sure the impact of bagging wasn’t being driven by motion, and so mean framewise displacement was regressed out from each individual mean adjacency matrix and ICC, MSE, and MSR were calculated. As was the case without motion regression, here we see that bagging improves the ICC, and reduces the MSE and MSR of parcellations (Supplementary Materials 5.2.3).

The individual voxel-voxel ICCs were moderate, reflecting that each of the voxel-voxel relationships that make up the individual parcellations are not highly statistically reliable; however, it is still possible that when taken as a whole, the parcellation patterns in these adjacency matrices would be reliably distinct between individuals. We tested this using the multivariate metric of reliability called discriminability (Figure 15). This method allows us to consider the extent to which individual parcellation patterns are unique to that individual compared to the rest of the group. Discriminability of the parcellations for the standard clustering approach was heavily impacted by the amount of data acquired and was quite high for 25 and 50 minutes of data (25-minute mean = 0.761; 50-minute mean = 0.817). We found that discriminability was significantly increased through bagging but not for all lengths of data. It seems that the individual level signals were not strong enough in 3-15 minutes of data to improve the discriminability of parcellation through bagging in the HNU1 data (all p > 0.05); however, we did see significant improvements from 0 to 400 bootstraps for 20 minutes (Kruskal-Wallis = 7.18, df = 1, p-value < 0.01), 25 minutes (Kruskal-Wallis = 29.44, df = 1, p-value < 0.0001); and 50 minutes (Kruskal-Wallis = 24.58, df = 1, p-value < 0.0001), showing individual parcellations became even more highly discriminable for 20 (mean = 0.71), 25 (mean = 0.89), and 50 (mean = 0.92) minutes.

**Figure 15.**
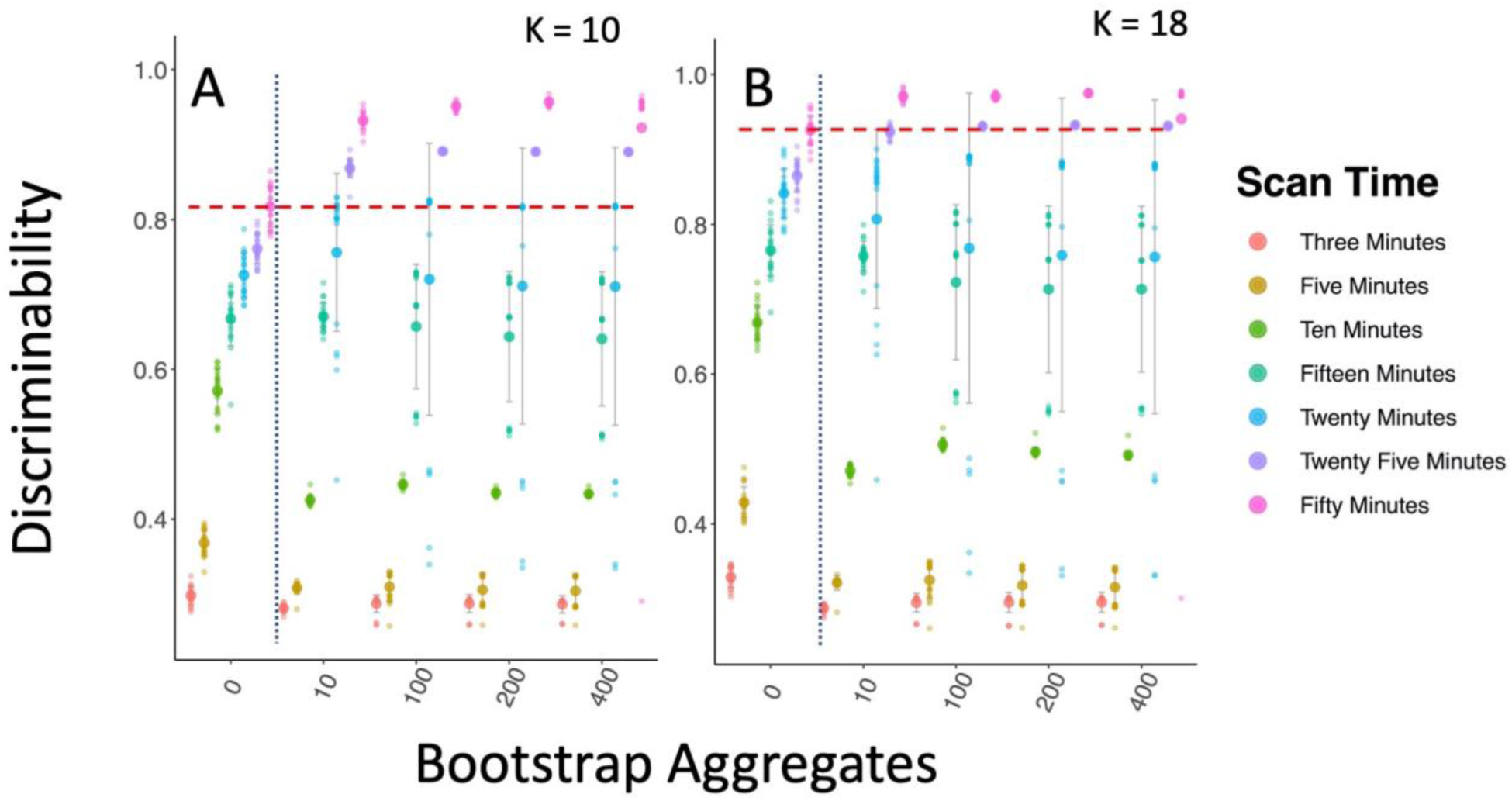
Bagging Improves Multivariate Reliability in Subcortical Parcellation. Figure A and B show the discriminability results for the subcortical parcellation for K = 10 and K = 18 respectively. The dotted blue line shows the standard clustering approach on the left, and the bagging approach on the right. The dashed red line shows the top mean performance for the non-bagged parcellations. Bagging leads to improved discriminability for longer scan times.

Finally, when making generalizations from group level information to the individual, it’s critically important that the group level information is representative of the individual (Adolf & Fried, 2019; Fisher, Medaglia, & Jeronimus, 2018). Such individual-to-group alignment is critical for deploying any group-defined models on individual level data, and as such is central for any kind of biomarker discovery process. Given that recent work has demonstrated that individual level brain networks can differ substantially from group level estimates (Gordon, Laumann, Adeyemo, & Petersen, 2015; Laumann et al., 2015), we wanted to test whether bagging could improve the individual-to-group similarity of parcellation, improving their potential for scientific and clinical applications (Figure 16). We found that bagging led to significantly more generalizable parcellations (20 minute parcellation; 0 vs 400 bootstraps; K = 10; Correlation: Kruskal-Wallis = 800.81, df = 1, p-value < 0.0001; ARI: Kruskal-Wallis = 248.43, df = 1, p-value < 0.0001), and increasing scan time did as well (K=10; Correlation: Kruskal-Wallis = 50.99, df = 1, p-value < 0.0001; ARI: Kruskal-Wallis = 16.59, df = 1, p-value < 0.0001). We also found that bagging led to an overall decrease in the variance of the individual-to-group similarity (K = 10; Correlation: F(599,599) = 0.56, p-value < 0.0001; ARI: F(1199,1199) = 0.85, p-value < 0.01), but this decreased variance was less pronounced here than in other analyses.

**Figure 16.**
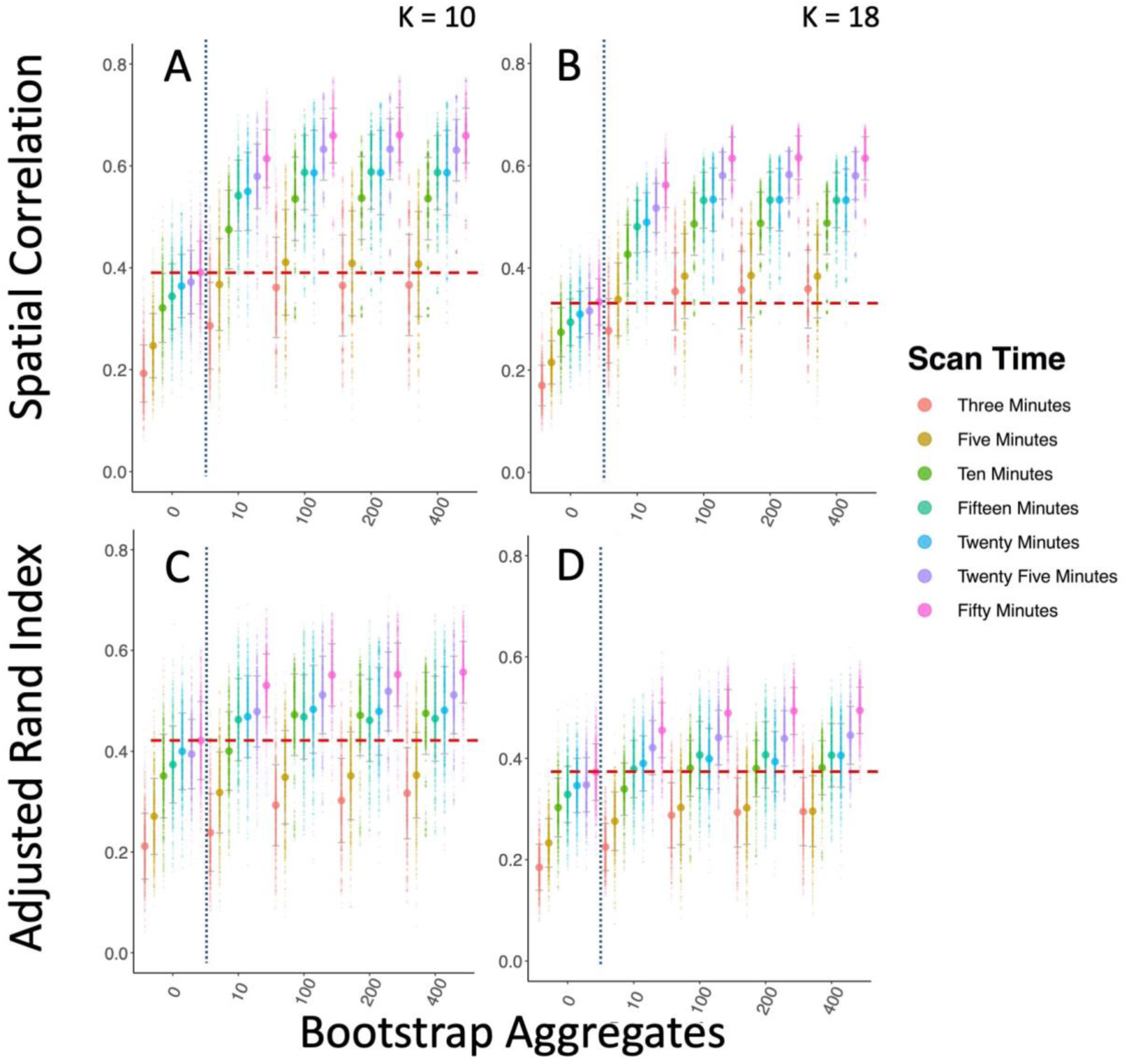
Bagging Increases Individual-to-Group Similarity of Parcellations. Overall this figure shows how bagging improves individual-to-group similarity between parcellations. The dotted blue line shows the standard clustering approach on the left, and the bagging approach on the right. The dashed red line shows the highest mean similarity of the standard parcellation approach. A & B show the improvement in correlation between the group-mean adjacency matrix and each individual’s mean adjacency matrix for the K = 10 and K = 18 subcortical parcellation respectively. C & D show the improvement of the ARI between the cluster’s labels of the group and individual level parcellations.

### 3.4 Characterizing Parcellations

Understanding how well parcellations map onto one another is an important element of assessing the likelihood of brain network results to generalize from one study to another. We assessed the extent to which our parcellations differed from one another and popular reference atlases. For example, we found that the differences between the HNU and GSP subcortical parcellations were most pronounced in the transition zones between parcels (Supplementary Figure 5.2.1), in particular we found the boundaries between the pallidum and putamen, and anterior superior thalamus to posterior inferior thalamus to be the source of greatest difference in the parcellations in the two datasets. Notably these differences were robust with both the 10 minute and 50-minute HNU data. Moreover, we replicate these areas of high differentiability between the HNU reference and replication datasets as well.

We also assessed the extent to which our subcortical parcellation aligned with previously established thalamic and striatal parcellations and we found that bagging neither improves nor reduces alignment between our subcortical parcellations and the FSL Harvard Oxford 3-cluster striatal solution and the FSL Harvard Oxford probabilistic 7-cluster thalamic parcellation (ARI: Kruskal-Wallis = 2.40, df = 4, p-value > 0.05). We also assessed the extent to which our cortical parcellations aligned with the Yeo and BASC atlases for K = 7. We found that bagging improved alignment for the Yeo atlas (ARI: Kruskal-Wallis = 5.86, df = 1, p-value < 0.02, but not for the BASC atlas (ARI: Kruskal-Wallis = 0.14, df = 1, p-value > 0.05). Notably our parcellations aligned with these atlases at level similar (ARI 0.2-0.3) to what would be expected comparing a wide range of atlases with different masks, samples, clustering parameters, variability in volume/surface alignment, etc., as parcellations with high similarity typically share many of these attributes (Myers et al., 2019). To understand the parcel distribution across the cortex and subcortical regions we also assessed the laterality of our parcels. The laterality of a parcellation is partially a function of the source of variance of the parcellation. For example, in some cases subcortical parcellations are conducted based on their connectivity to bilateral cortical networks (Choi et al., 2012), in which case the parcels may be mostly or exclusively bilateral, because the parcellation is segregating them with respect to their association to a relatively symmetric signals across hemispheres of cortex (Yeo et al., 2011; Bellec et al., 2015). In other cases, such as those when the subcortical areas are parcellated based on the similarity of within-region time series (Kim, Park, & Park, 2013), differences between right and left hemispheres may result in lateral differences in the parcellation, as is the case in our subcortical parcellation here. We found that over 20 repeated parcellations of the subcortical regions (20-minute HNU replication data; K = 10) about half of the parcels spanned both hemispheres (40.4%), aligning with prior work (Kim, Park, & Park, 2013). We also found that in 10 repeated parcellations of the cortex (10-minute HNU replication data; K = 7) most of the parcels were bilateral (82.9%), which aligns with previous results as well (Yeo 2011, Bellec et al., 2015). Notably we found that bootstrap aggregation did not change the number of bilateral parcels in either parcellation scheme (Subcortical: Kruskal-Wallis = 7.98, df = 4, p-value > 0.05; Cortical: Kruskal-Wallis = 0.0089, df = 1, p-value > 0.05).

## 4. Discussion

### 4.1 Overview

Our analyses revealed that bagging improves the between sample reproducibility of group level functional parcellations, and that this effect generalizes across multiple subsamples, clustering parameters, and clustering algorithms. Furthermore, we replicate these effects in both cortical and subcortical parcellations. In many cases, bagging improved reproducibility more than doubling scan time. As such, bagging appears to be a valuable means of bringing down data requirements for the reproducible study of functional brain organization. Bagging enhanced parcellations were also more reproducible and reliable at the individual level, with an order of magnitude decrease in the MSE and MSR of individual level parcellations and significant increases in univariate and multivariate reliability. With twenty or more minutes of data we saw increases in discriminability for individual level parcellations that were already highly discriminable. We saw bagging driven increases in between session individual level reproducibility across all scan lengths and both spatial correlation and ARI metrics. This implies that the highly detailed, individual-specific attributes of parcellations can be further enhanced through bagging. We found that higher parcel number was associated with lower group level reproducibility, and we found bagging alleviated some of this decrease in reproducibility, with more detailed bagged parcellations (K = 197) being as reproducible as less detailed, network-level non-bagged parcellations (K = 7). This suggests that bagging may be especially useful for achieving acceptable reproducibility when moving towards finer spatial scales. We found that bagging enhanced parcellations also had higher between session and between scan reproducibility. This suggests that bagging enhanced parcellations are better suited for within-sample repeated measures that are commonly used in intervention or other longitudinal studies in clinical and cognitive neuroscience. Critically, this also demonstrates that our results generalize from low resolution “network-level” clustering to the more detailed “parcel-level” clustering with hundreds of regions. Finally, we found that bagging improved individual-to-group similarity of parcellations, which is key for minimizing parcellation mismatch and as a result, the potential for artifactual findings. Overall, our findings indicate that bagging enhanced parcellations: 1) outperform the standard approach on a wide variety of reproducibility and reliability indices, and 2) should be considered for further implementation in cutting-edge cluster and gradient-based parcellation and other approaches for potential improvements in making more robust measurements of the connectome (Glasser et al., 2016; Laumann et al., 2015; Schaefer, Kong, & Yeo, 2016; Xu et al., 2016).

### 4.2 Why Bagging Works

Both supervised and unsupervised machine learning methods suffer from variability of estimation that are inherent to aspects of the models themselves (Tumer & Ghosh 1999). For example, K-means is known to be unstable and sensitive to outliers (Li et al., 2011). Furthermore, when working in high dimensional spaces, as is often the case with clustering, variability in model estimation also arises due to the curse of dimensionality (Friedman, Hastie, & Tibshirani, 2001). As distances are computed in more dimensions, the density of sampling within each dimension also decreases. This sparsity of observations per dimension leads to increased variability in model estimation for a given sample or across samples. This variability in sampling means that relatively small perturbations in the either the sample, distance metrics used by the clustering algorithm, or clustering parameters, can lead to significant variability in the cluster outcomes. Detailed review of different parcellation techniques shows that while none are necessarily superior across all conditions, they can be quite different from one another (Arslan et al., 2018; Kleinberg, 2003; Priebe et al., 2019).

To overcome this variability, aggregating over multiple resampled datasets can provide a more generalizable model. When assessing a range of models, each individual model captures different elements of the structure of the data. Even if the best model were selected, important information contained in the results of the discarded, “non-optimal” models may be lost. Bagging enables the final decision to be made using information from all models collected across a range of resampled datasets, thereby mitigating this loss of information (Tumer & Ghosh, 1999). This effect of model aggregation has been well understood in the supervised learning literature (Breiman, 2001; Krogh & Vedelsby, 1995; Tumer & Ghosh, 1999), and has more recently been shown to provide similar benefits in unsupervised domains as well (Hu & Yoo, 2004; Lancichinetti & Fortunato, 2012; Strehl & Ghosh, 2002). In order for the aggregated models to improve performance, they must be different from one another (Boongoen & Iam-On, 2018; Krogh & Vedelsby, 1995; Kuncheva & Hadjitodorov, 2004), and the diversity of the aggregated models is linked to the benefits seen through aggregation (Breiman, 2001; Hadjitodorov, Kuncheva, & Todorova, 2006). By using a multilevel bagging approach in the current work, we are able to capture information from all the variability within individuals and across individuals. We see that both levels contribute to improved parcellation reproducibility due to their ability to aggregate over different non-overlapping sources of variance.

Beyond nuisance signals (e.g., head motion, respiration, cardiac variation), the largest source of within-individual variance likely comes from the dynamic nature of functional connectivity (Chang & Glover, 2010; Hutchison et al., 2013). One possibility is that bagging may be improving parcellation stability by making the final clustering representative of the most dominant and stable trends in the data. With a clearer model of the dominant or central tendency of connectivity patterns in a brain region, bagging seems to prevent small perturbations in the data from having outsized effects on the parcellation results. We believe this contributes to the ability of bagging to improve parcellation reproducibility across scans, sessions, samples, and scanners.

Variability between parcellations also arises due to specific biases inherent within each clustering algorithm. While the field has not established any ‘best’ technique for functional parcellation (Arslan et al., 2018) or clustering in general (Kleinberg 2003; Priebe et al. 2019), bagging does not rely on any particular clustering technique. Though we demonstrate the impact of bagging in clustering here, bagging can be also applied in a wide range of other functional parcellation approaches (i.e. multivariate decomposition, gradient-based parcellation). With continuing advances in functional and multi-modal parcellations, bagging can be employed in new methodological approaches to continually improve parcellation reproducibility, reliability, and individual-to-group similarity. There may also be other ways in which ensemble methods and bagging can be used to improve the generalizability of functional parcellations and other MRI approaches that use clustering (Blumensath et al., 2013; Craddock et al., 2012). For example, prior work has demonstrated the utility of using clustering for tissue class segmentation in lesion detection (Bosc et al., 2003; Juang & Wu, 2010); these efforts may be furthered by the use of cluster ensemble methods for detecting and evaluating differences lesions more reproducibly across samples and timepoints.

### 4.3 Improving Reproducibility

Within the scope of human neuroscience, the reproducibility of functional parcellations is especially important. Parcellations have a significant impact on the sensitivity of the connectome to phenotypic variables (Abraham et al., 2017), and there are key issues that arise when parcellations vary in their fit across groups and individuals. For example, recent work has demonstrated that up to 62% of the variance in the edges of a connectome can be explained by the alignment of the reference parcellation to the data in question (Bijsterbosch et al., 2018). This is troublesome because when parcellations do not generalize well between samples, the resulting connectome edges will conflate differences between group network structure with differences in underlying parcellation fit (Zuo & Xing, 2014). This leads to purported markers of psychiatric illness failing to reproduce, and therefore limiting production of biomarkers in psychiatry. When considering failures to replicate (e.g., (Dinga et al., 2018; Drysdale et al., 2017), it is important to consider potential limitations in the generalizability of the template parcellation applied across samples. There are a multi-faceted set of study-related differences that may exist between studies that have created the most commonly used brain atlases. This includes the differences in the kind of data acquired, including number of participants, the length of MRI scans, acquisition parameters, and scanner type. More generally, the composition of study samples may differ considerably on a range of phenotypic variables including age, sex, socioeconomic status, cognition and psychopathology (Falk et al., 2013). When parcellations are not sufficiently generalizable, it becomes more difficult to assess whether differences in functional connectivity between individuals or groups are due to inherent differences in connectivity strength between areas, or differential fit of a parcellation. In this way, reproducible parcellations are essential for creating reproducible relationships between functional connectivity estimates and behavioral, cognitive, or clinical variables. Our finding that bagging improves parcellation reproducibility at both the group and individual level suggests bagging may be a key element to creating maps of functional organization that allow for robust estimations of brain-behavior relationships.

### 4.4 Lowering Data Requirements

We found that bagging enhanced parcellations allowed us to lower our minimum data requirements for generalizable functional parcellations. There are many efforts that have demonstrated that robustness and generalizability of functional neuroimaging measurement are increased considerably by increasing the length of the fMRI scan to 25 or more minutes (Elliott et al., 2019; Laumann et al., 2015; Xu et al., 2016). However, this length of time may not be appropriate for some clinical communities, as long scan times are not feasible in many developing or patient populations. Furthermore, many currently existing datasets in clinical populations have between 5-10 minutes of scan time, such as the 1000 Functional Connectomes Project (http://fcon_1000.projects.nitrc.org/fcpClassic/FcpTable.html), GSP, ABIDE I, ABIDE II and PING datasets (Di Martino et al., 2017; Di Martino et al., 2014; Jernigan et al., 2016). Therefore, finding methods to increase the stability of functional imaging without increasing scan time may assist in providing datasets of clinical populations more appropriate parcellations for their shorter scan times. Our results suggest that bagging may be a useful technique for creating more generalizable functional parcellations with less data.

### 4.5 Better Individual-to-Group Generalization

Bagging enhanced parcellations also have greater individual-to-group similarity compared to the standard parcellation procedure. Individual-to-group generalization is key for any scientific models attempting to make individual level inferences based on group level models or estimations. Given that an individual system may be unique in ways that share little relationship to group models (Fisher et al., 2018), and extensive individual-variation has been demonstrated in functional networks (Laumann et al., 2015; Xu et al., 2016), it’s also critical that if group models are used, they are as representative of each individual in the group as possible. In other words, individual-to-group representativeness is critical to make accurate assessments of individual level functional connectivity. Otherwise, within group differences between individuals in edgewise connectivity will also be the result of the parcellation fit, rather than actually the functional connection of interest (Bijsterbosch, Beckmann, Woolrich, Smith, & Harrison, 2019; Bijsterbosch et al., 2018). This brings the validity of the resulting brain-behavior relationships into question, as the maps being compared are assumed to be equivalent but are not mapping the territory equally well. Our results demonstrate that bagging may at least partially alleviate these issues by increasing the similarity of individual-to-group parcellations, thereby leading to more valid brain-behavior estimation.

### 4.6 Improved Individual Level Parcellations

We find that bagging improves reproducibility of individual level parcellations. With extensive recent efforts focusing on the importance of developing robust parcellations at the individual level (Gordon et al. 2015; Salehi et al. 2020; Xu et al. 2016), this finding is particularly important for improving estimation of brain-derived individual differences. Our findings demonstrate that bagging leads to a ten-fold decrease in the variance of individual level parcellations between sessions, and a nearly ten-fold decrease in the variance of parcellations between individuals as well. Decreasing individual variance in parcellation over time implies that bagging enhanced parcellations may be more able to replicate individual level parcellation patterns in the pre sence of noise. Lower variance between individuals suggests that bagging enhanced parcellations bring individuals closer together, potentially removing irrelevant differences between individuals in parcellation patterns. This interpretation is supported by our findings showing that with enough scan time, bagging enhanced parcellations also produce parcellations that are more discriminable, meaning that the individual-specific parcellation patterns are enhanced. Discriminability is a key criterion for distinguishing between individuals and groups on the basis of phenotypic differences. Developing methods that enhance the discriminability of existing data enables greater sensitivity and understanding of brain-behavior relationships. By making parcellations that more accurately represent the individual, it may be possible that bagging enhanced parcellations also result in functional network edges and other metrics that are more reliable, discriminable, and are more sensitive to brain-behavior relationships as well.

### 4.7 Improved Within Sample Repeated Measures

Assessing change over time relies on within sample reproducibility, at both the individual and group levels. Studies of brain development create accurate phenotype-brain relationships by using large samples to overcome variability due to measurement error (Volkow et al., 2018). However, for cognitive or clinical interventions studies (Anguera et al., 2013; Nikolaidis, Voss, Lee, Vo, & Kramer, 2014; Oelhafen et al., 2013), collecting such large samples is completely unfeasible. Therefore, creating methods that improve the between session reproducibility of neuroimaging measurements may increase sensitivity of analysis to intervention related changes. By decreasing variance between individuals, within individuals, and across repeated measures, we create parcellations that may be better suited to pick up individual differences in functional connectivity that are driven by an effect of interest. This is an essential point for improving reproducibility of functional maps of the brain and bringing functional MRI closer to clinical impact.

### 4.8 Limitations

There are a large range of optimizations that may further improve parcellation reproducibility that were not investigated in the current work. For example, there are several preprocessing choices that may impact the reproducibility of parcellations, such as registration optimization, denoising parameter selection, and exclusion of global signal regression. While examinations of these optimizations are outside the scope of the present work, they may have important interactions with bagging that could have an impact on the parcellation improvements we see here. We also note that our parcellations did not align very highly with established subcortical and cortical parcellations. It’s important to note that different techniques all produce inherently different parcellation schemes due to differences in surface/volume brain alignment, cluster number, regions of interest used, and cluster parameters. Furthermore, the current work has noteworthy limitations on the samples chosen. We only used healthy young adults in both samples; considering how to deal with the presence of meaningful between individual differences (e.g., development status, clinical status) when bagging in more heterogeneous samples requires further work. Another limitation of the current study is that we did not exhaustively test the space of cluster methods, which are known to vary in their reproducibility (Arslan et al., 2018), even though this is possible with PyBASC (See Supplementary Materials 5.1). Bagging and cluster ensembles are notable for being agnostic to clustering technique and have been successfully demonstrated in a wide range of clustering techniques however, and we feel this limitation is notable but not a significant one. Finally, the bagging procedures executed in the current work add computational requirements to the creation of parcellations. Therefore, when very large datasets or very high-resolution functional data are used, this technique may not be technically feasible. This may change in the future as more advanced computational resources become available. There are a range of other optimizations yet to be investigated, and the current work is just one step in that direction.

### 4.9 Future Directions

In the age of the ‘replication crisis’ in neuroimaging, methods to improve the reproducibility of our neuroimaging measurement are absolutely essential. Bagging for improved cluster stability is not a new technique, nevertheless its rarely used in functional parcellation and clustering in neuroimaging. We hope that the current effort has addressed a lack in the extant literature on the benefits of bagging on between sample and between session reproducibility of cluster estimation. While the current effort aimed at establishing the role of bagging in improving reproducibility and reliability of functional parcellations, the number of tests were far from exhaustive. There are more ways that bagging and cluster ensembles could be leveraged to improve measurements of the brain. For instance, varying and aggregating over a range of time series window lengths, cluster techniques, or resampling methods may lead to better individual level and group level estimates. Some research has demonstrated that selecting diverse sets of cluster solutions for aggregation can outperform simply aggregating all cluster solutions (Hadjitodorov et al., 2006). Such a diversity selection approach could also be employed in a similar context and may likely improve reproducibility and reliability of the cluster solutions as well. Given that bagging improves these qualities in parcellations, it’s possible that bagged parcellations may also show improved reproducibility and reliability of edgewise functional connectivity. Furthermore, these edges may also be more sensitive to phenotypic differences between people and groups, and these considerations should be explored in future work.

## Acknowledgements

We would like to thank Lei Ai and Yu Tong for their assistance with the GSP and HNU1 datasets. We would also like to thank the extensive open science infrastructure that supported the creation of this work, in particular the OHBM and Brainhack communities for their input and support during the creation of this project. We would like to thank Xinian Zuo and Randy Buckner and the teams behind the creation and release of the HNU1 and GSP datasets, as well as the participants involved in both. We would also like to thank the teams involved in the creation of Nilearn and Scikit Learn, two programming resources that have been invaluable to the current work.

## 5. Supplementary Materials

### 5.1 Supplementary Methods

#### 5.1.1 Standard Clustering Resampling Datasets

Without bagging, Ward’s criterion hierarchical agglomerative cluster is deterministic and has no variance over multiple runs. To offer a better understanding of the sampling variance that could lead to differences in clustering when bagging is not applied, we apply resampling to the original replication datasets when comparing them to a given reference. To create the multiple 3- and 5-minute datasets for the non-bootstrapped condition, we randomly sampled continuous 3- or 5-minute blocks of the full 50 minute replication dataset. To create the 10, 15, 20, 25, and 50-minute resampled datasets, we randomly sampled continuous 5-minute blocks of the full 50-minute replication dataset. We did not calculate resampling in the GSP datasets because we compared the 10 groups against one another.

#### 5.1.2 PyBASC

Originally, this method was implemented in Octave, and we have recently implemented this method in Python to take advantage of the broad range of powerful libraries available for machine learning and neuroimaging, including NiBabel, Nilearn, Scikit Learn, and NiPype. This enables PyBASC to be highly flexible and extensible in adopting new clustering algorithms and implementations, but more importantly allows for dramatic increases in computational power through the use of NiPype’s easy-to-use and adjust multi-core processing capabilities. We have also made edits to the processing that make PyBASC run very efficiently with minimal memory loads. This makes PyBASC just as easy to run on a laptop as on a large server in the cloud. Here, we briefly address the PyBASC overall methodology. For more information, we have also explained these methods in more detail elsewhere (Bellec et al., 2010; Garcia-Garcia et al., 2017). PyBASC has a wide range of functionalities that can be customized according to user preferences. Well-performing default values are provided where applicable, but users can choose 1) the number of initial supervoxels, 2) the number of individual and 3) group-level bootstraps, the 4) region to be parcellated and 5) the secondary region (optional) to use as the basis of cross-network clustering. Users can also define the cluster method to be used, as well as the affinity threshold and distance metric used in the clustering procedures.

#### 5.1.3 Individual-Level

First to reduce the dimensionality of the voxel wise time series, and to allow for better computational efficiency, we apply a hierarchical feature agglomeration from Scikit Learn with a neighboring voxel spatial constraint to each individual’s data (Subcortical Parcellation: 600 supervoxels; Cortical Parcellation: 1800 supervoxels). Then, a circular block bootstrap procedure is applied to create resampled time series data from the original functional image. We follow prior recommendations to set the block length to greater than 20 time steps (Adolf et al., 2014), and more specifically, equal to the square root of the number of points in a time series (Bellec et al., 2010; Bellec et al., 2008). Each of these resampled time series are then clustered using the clustering technique of choice. Currently K-means, Ward’s criterion hierarchical agglomerative, spectral clustering, and Gaussian mixture models are implemented. With each application of clustering, the cluster labels are transformed into an adjacency matrix of 0s and 1s representing whether a given voxel belongs to the same cluster as another voxel. Each of these adjacency matrices are in supervoxel space, which differs between individuals. Therefore, the dimension reduction procedure is reversed to create a voxel wise adjacency matrix that puts all individual level data back into the same space. All adjacency matrices are averaged together to create an individual mean adjacency matrix (IMA), which represents a summary of all clustering assignments across all bootstraps. The IMA can be clustered to produce a final consensus cluster assignment for each individual, and it can be used in group level bagging.

#### 5.1.4 Group Level

PyBASC applies bagging to IMAs across individuals, recreating resampled groups of IMAs from the original pool of participants. The IMAs for each group level bootstrap are averaged and another group adjacency matrix is created through clustering. The adjacency matrix across all bootstraps are averaged to create a group mean adjacency matrix (GMA), and this GMA is clustered to create the final group level clustering. The GMA is the final output of the group level bagging, along with the cluster labels. While the IMA represents a summary of the clustering across all bootstraps for each individual, the GMA represents a summary of the clustering across all bootstraps for the group. We compare each individual’s IMA to the GMA as a detailed way of assessing similarities between the individual and group level clustering.

### 5.2 Supplementary Results

**5.2.1.**
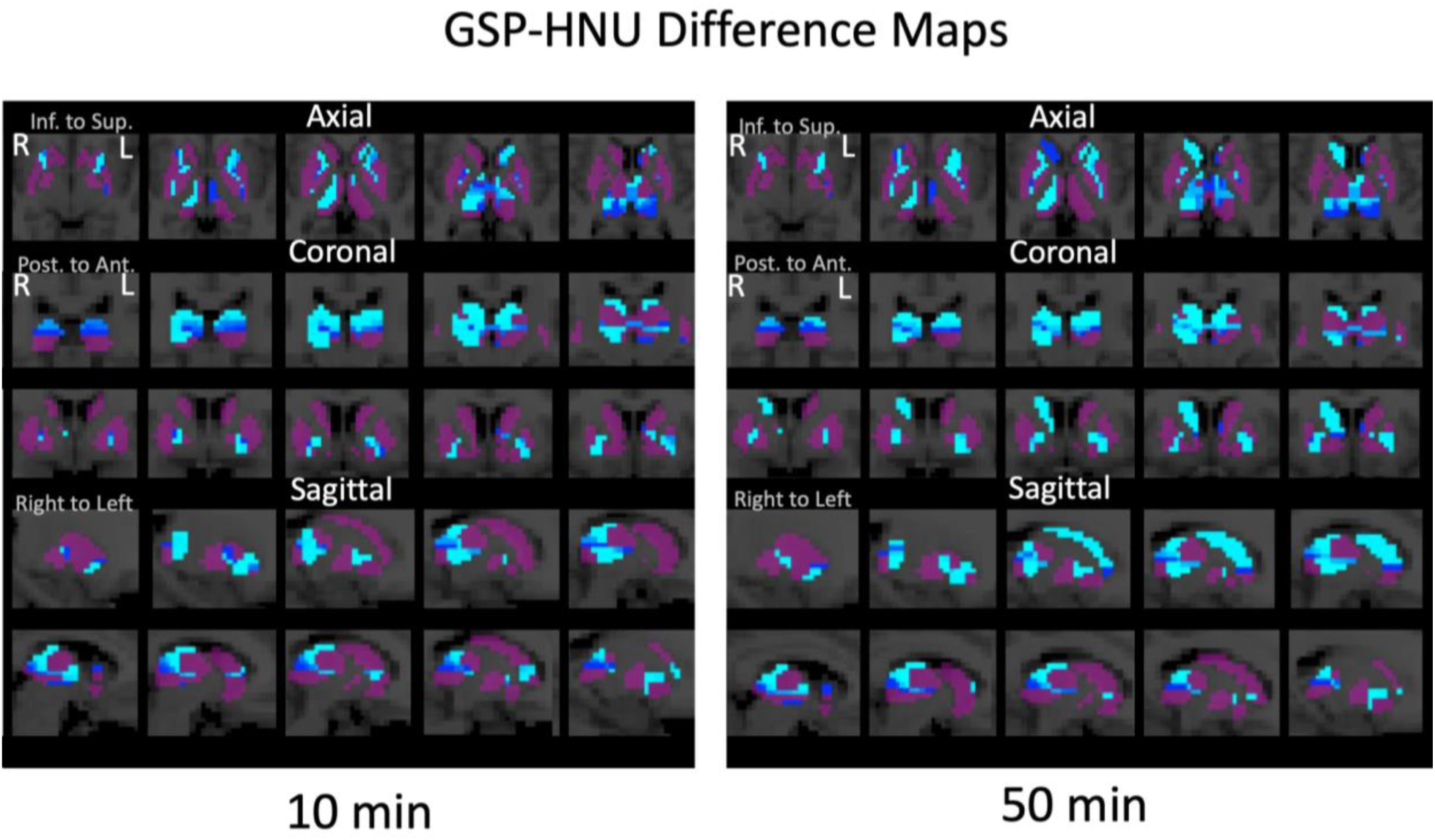
Here we compare the voxelwise cluster assignment differences between the Reference GSP dataset created with 400 bootstraps and the 10- and 50-minute HNU1 parcellations. Bright blue corresponds to all comparisons being different between the two parcellations across all replication samples with 400 bootstraps, and dark purple shows where there are no differences between parcellations. We find that both 10- and 50-minute parcellations mostly differ from the GSP in transition zones between parcels. In particular we see these difference maps reveal a superior-inferior thalamic transition zone, and a putamen-pallidum transition zone. Notably, the HNU1 50-minute parcellation differs from the GSP reference in differential classification of the caudate. Whereas the 50-minute HNU1 data clusters the bilateral caudate into the same cluster, both the GSP and the 10-minute HNU1 data cluster the caudate into separate, non-homologous clusters.

**5.2.2.**
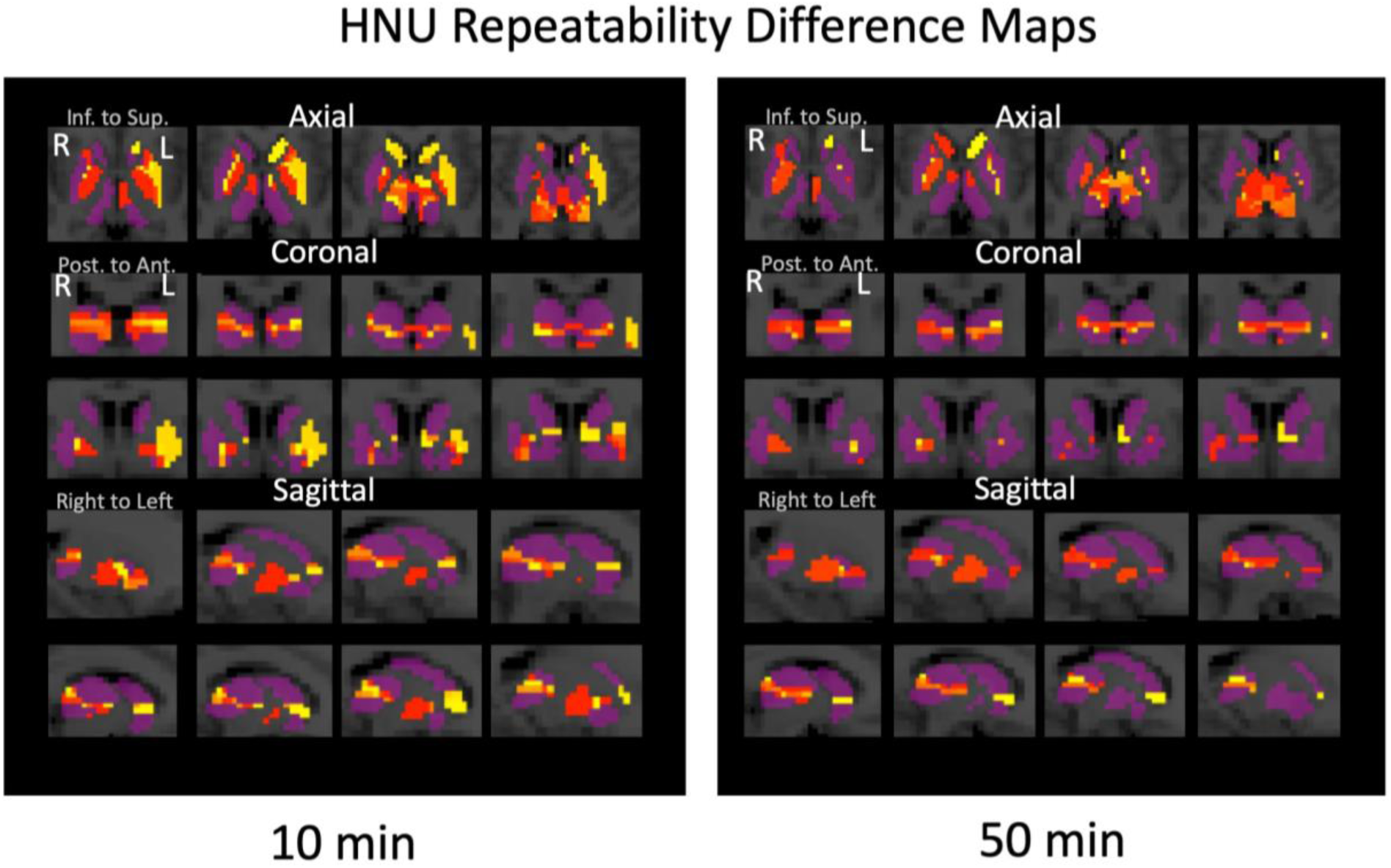
Here we compare differences the HNU1 Replication and Reference parcellations. Bright yellow corresponds to all comparisons being different between the two parcellations across all replication samples with 400 bootstraps, and dark purple shows where there are no differences between parcellations. Notably we see differentiation between reference and the 10-minute parcellation is higher than the 50-minute replication set. Furthermore, we see a reprise of the transition zones seen in S1, notably a clear superior-inferior transition, and a putamen-pallidum transition as well.

**5.2.3.**
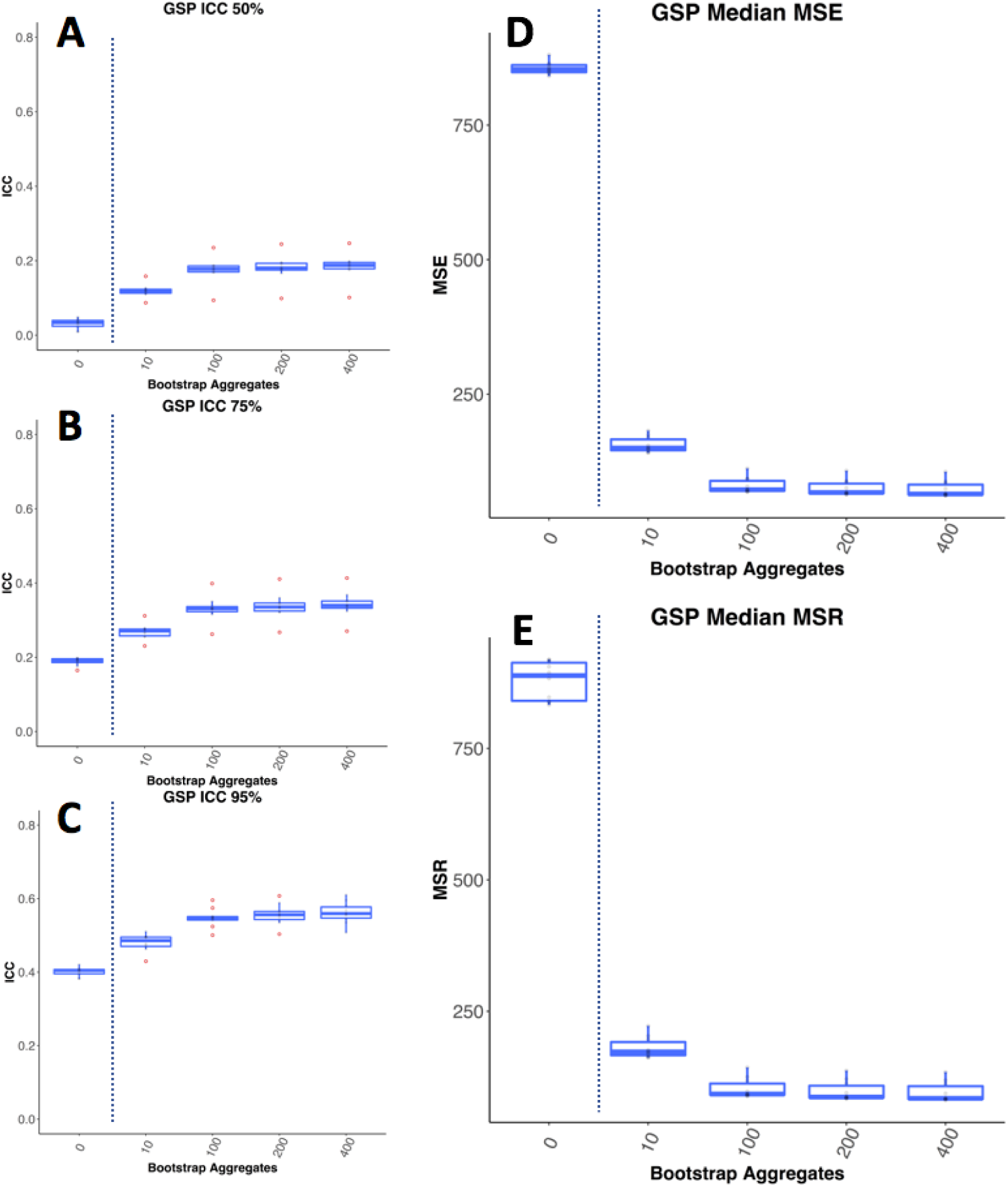
Univariate Reliability in GSP with motion regressed out. Motion can be a source of highly reproducible signal in MRI data. We wanted to make sure the impact of bagging wasn’t being driven by motion, and so we regressed out motion from each individual mean adjacency matrix before calculating ICC. As was the case without motion regression, here we see that bagging improves the ICC of parcellations. A,B,C show the top 50%, 75%, and 95% ICC values for the individual level mean adjacency matrices. Each point corresponds to the 50th, 75th, and 95th percentile ICC values for the individual level mean adjacency matrices for each of the 10 GSP groups. D and E show the mean squared error within individuals and between individuals respectively. We see that bagging improves ICC, it has a dramatic impact on both MSE and MSR, with samples with the lowest drop in MSR showing the greatest pickup in ICC.

**5.2.4.**
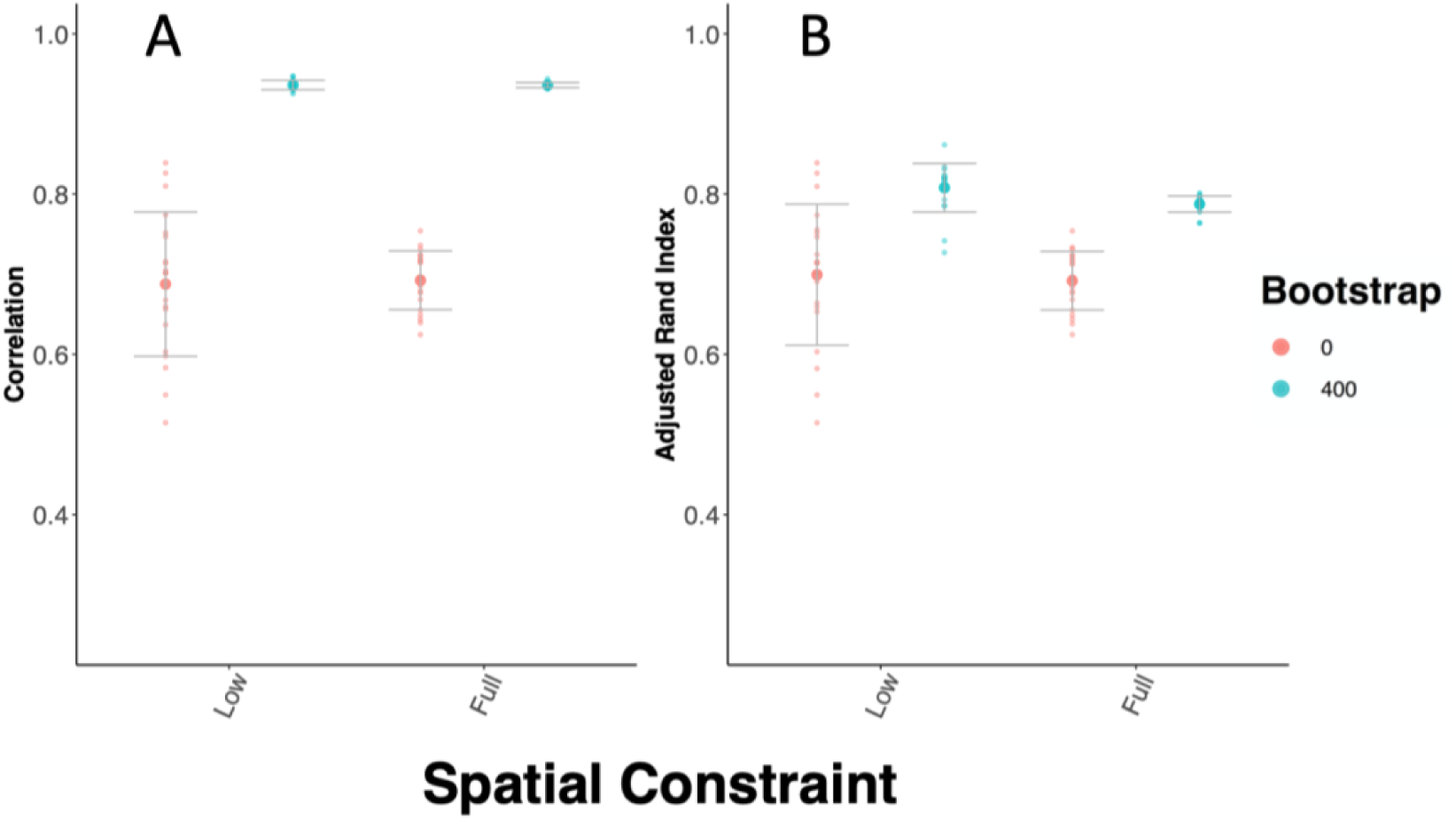
<u>Bagging improvements generalize across spatial constraints.</u> This figure demonstrates bagging improves between session reproducibility in subcortical parcellation across levels of spatial constraints. In the ‘Low’ spatial constraint condition, the approximately 1800 voxels of the subcortical region of interest are reduced slightly to 600 supervoxels separately for each individual. In the “Full” spatial constraint condition, the subcortical parcels are fully spatially constrained, meaning that no discontiguous parcels exist. We find that bagging improves reproducibility of the parcellations in both these conditions.

**5.2.5.**
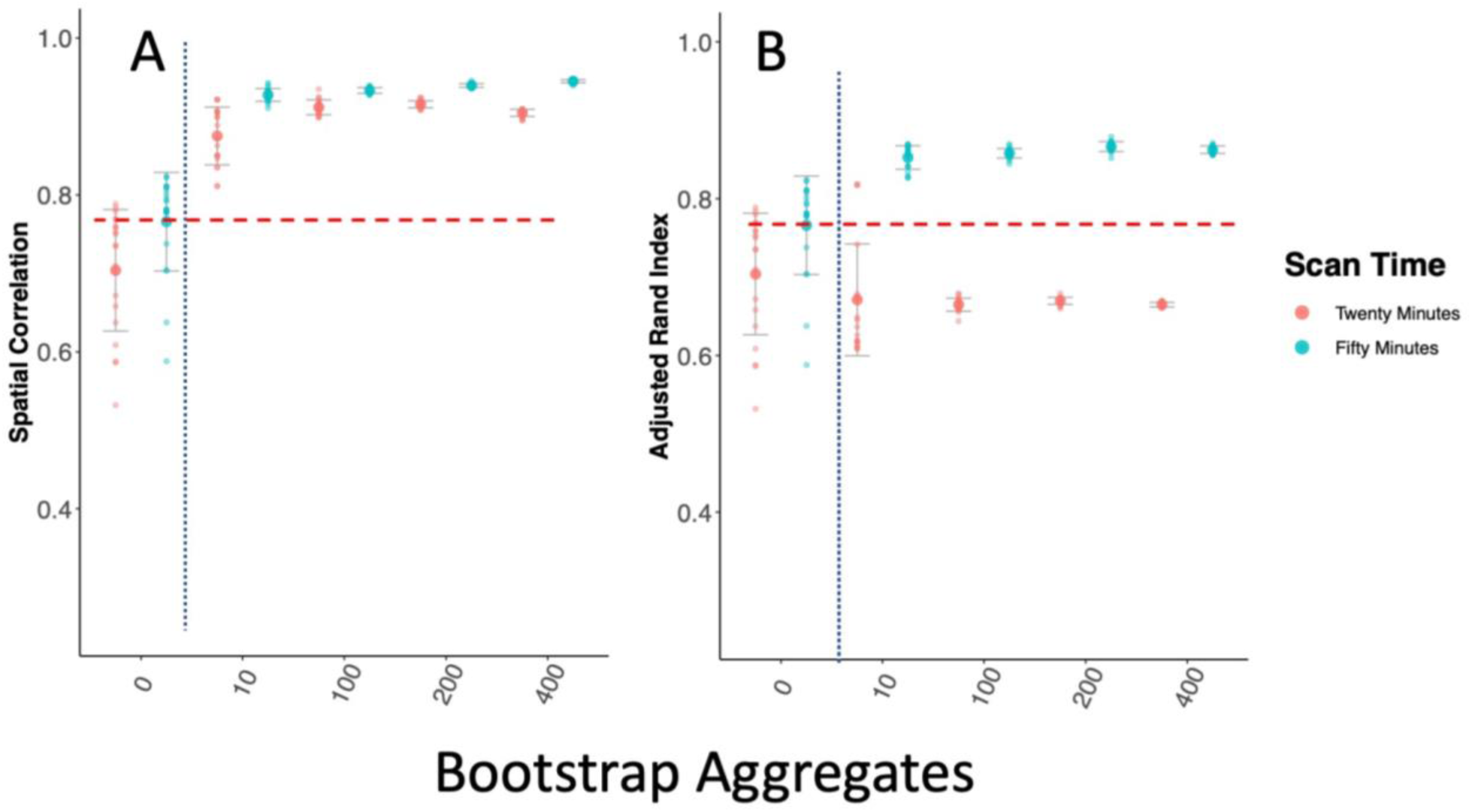
<u>Bagging Improves between session reproducibility of subcortical parcellation with spectral clustering.</u> This figure demonstrates that the beneficial impact of bagging generalizes across clustering techniques as well. Notably however, while the spatial correlation of the parcellation improves considerably with bagging for both 20- and 50-minute scans, ARI only shows significant improvements for 50 minutes of data.

